# Neuron secreted chemokine-like Orion is involved in the transformation of glial cells into phagocytes in different neuronal remodeling paradigms

**DOI:** 10.1101/2022.12.16.520775

**Authors:** Clarisse Perron, Pascal Carme, Arnau Llobet Rosell, Eva Minnaert, Salomé Ruiz Demoulin, Héloïse Szczkowski, Lukas Jakob Neukomm, Jean-Maurice Dura, Ana Boulanger

## Abstract

During animal development, neurons often form exuberant or incorrect axons and dendrites at early stages, followed by the refinement of neuronal circuits at late stages. Neural circuit refinement leads to the production of large amounts of neuronal debris in the form of neuronal cell corpses, fragmented axons and dendrites, and pruned synapses requiring disposal. In particular, the predominant phagocytes acting during the neuronal remodeling and degeneration are glial cells and critical signaling pathways between neurons and glia leading to phagocytosis are required. Chemokine-like mushroom body neuron secreted Orion ligand was shown to be essential to the astrocyte infiltration into the γ axon bundle leading to γ axon pruning and clearance of debris left from axon fragmentation. Here we show a role of *orion* also in debris engulfment and phagocytosis. Interestingly, we show that *orion* is also involved in the overall transformation of astrocytes into phagocytes. In addition, analysis of several neuronal paradigms demonstrates the role of *orion* in the elimination of both peptidergic vCrz^+^ and PDF-Tri neurons *via* additional phagocytic glial cells as cortex and/or ensheathing glia. Our results suggest that Orion is essential for phagocytic activation of three different types of glial cells: astrocytes, cortex and ensheathing glia and point to Orion as a trigger not only of glial infiltration but also engulfment and phagocytosis.

## Introduction

During animal development, neurons often form exuberant or incorrect axons and dendrites at early stages, followed by the refinement of neuronal circuits at late stages. Neural circuit refinement can proceed by the complete elimination of a neuron and its projections or the selective destruction of specific axons, dendrites, or synapses. During development, this elimination occurs by different mechanisms like: axonal degeneration, axon retraction or cell apoptosis. Similar molecular and cellular mechanisms are at work during neurodevelopmental disorders or after nervous system injury (Luo and O’Leary 2005; Neukomm and Freeman 2014; Riccomagno and Kolodkin 2015; Schuldiner and Yaron 2015; Llobet Rosell and Neukomm 2019).

Neural circuit refinement leads to the production of large amounts of neuronal debris in the form of neuronal cell corpses, fragmented axons and dendrites, and pruned synapses requiring disposal. In particular, the predominant phagocytes acting during the neuronal remodeling and degeneration are glial cells and critical signaling pathways between glia and neurons leading to phagocytosis have been recently identified (Boulanger and Dura 2022). The elimination of neuronal debris by glial cells can be divided into three different cellular steps. The first step is the infiltration of axon bundles by glial cells to reach the target to be degraded. The second is the recognition/engulfment of neuronal debris. The third step is the phagocytosis through the endocytosis of the engulfed debris by glial cells resulting in the formation of phagosomes which subsequently mature.

In *Drosophila*, the mushroom body (MB), a brain memory center, is remodeled at metamorphosis and MB γ neuron pruning occurs by a degenerative mechanism (Watts et al. 2003; Yu and Schuldiner 2014; Boulanger and Dura 2015; Yaniv and Schuldiner 2016). The γ axon fragmentation is under the control of the MB intrinsic γ neuron program which depends essentially on the ecdysone receptor whose transcriptional regulation is finely orchestrated (Boulanger et al. 2011; Boulanger and Dura 2015). Astrocytic glia surrounding the MB have an active role in the process; blocking their infiltration into the MBs prevents remodeling (Awasaki and Ito 2004; Hakim et al. 2014; Tasdemir-Yilmaz and Freeman 2014). Thus, infiltration of astrocytic processes into the MB γ axon bundle seems to be an essential first step for the elimination of these axons by glia. In the ventral nerve cord (VNC), astrocytes already infiltrate the neuropil early in development, and transform into phagocytes removing excess of synaptic terminals at the initiation of metamorphosis. Elimination of peptidergic VNC Corazonin expressing neurons (vCrz) is a known example of VNC neurons whose neurites are eliminated by astrocytes. Interestingly, the cell bodies of these neurons were cleared by non-astrocyte glia (Tasdemir-Yilmaz and Freeman 2014). Thus, even though astrocytes have been identified as the major phagocytic cell type responsible for engulfing degenerating axons (Doherty et al. 2009; Stork et al. 2014) other glial cell-types as: ensheathing glia, cortex glia, and wrapping glia are involved in remodeling (Bittern et al. 2021; Boulanger and Dura 2022). These phagocytic glial cells are located in different regions of the central and peripheral nervous system. Recent studies have shown that peptide-dispersing factor tritocerebrum (PDF-Tri) neurons die by apoptosis following adult eclosion (Gatto and Broadie 2011) and are eliminated by ensheathing and cortex glia but not by astrocytes (Vita et al. 2021). In addition, ensheathing glia has a major role in the elimination of injury induced debris. Several examples of the ensheathing glia function on olfactory receptor neuron (ORN) debris phagocytosis after antenna or palp ablation have been reported (Doherty et al. 2009; Musashe et al. 2016). The role of wrapping glia in neuronal remodeling was poorly illustrated even though altering wrapping glia function blocks NMJ remodeling during metamorphosis (Boulanger et al. 2012). Moreover, after section of the L1 wing nerve, expression of the phagocytosis receptor Drpr (*draper*) in wrapping glia was essential for the elimination of the resulting debris, suggesting a phagocytic role of this type of glia in wing nerve after injury (Neukomm et al. 2014).

Little is known about the signaling pathways and neuron secreted ligand that activate glia and lead to their phagocytic transformation. Some ligands like phosphatidyl serine, Ilps, Spz, sAppl (Boulanger and Dura 2022) were shown to be secreted or presented by neurons to glia to activate phagocytosis pathways. We recently isolated the chemokine-like MB neuron secreted Orion ligand by EMS mutagenesis (Boulanger et al. 2021). Orion was shown to be essential to the astrocyte infiltration into the γ axon bundle leading to γ axon pruning. Moreover, we showed that the significant number of axonal debris seen in adult *orion* null individuals was due to the failure of astrocytes to clear debris left from axon fragmentation. Orion is a secreted protein and bears some chemokine features such as a CX3C motif and three glycosaminoglycan (GAG) binding consensus sequences that are required for its function. Chemokines are a family of chemoattractant cytokines characterized by a CC, CXC, or CX3C motif promoting the directional migration of cells. Mammalian CX3CL1 (also known as fractalkine) is involved in neuron-glia communication (Paolicelli et al. 2014; Arnoux and Audinat 2015; Wilton et al. 2019). Fractalkine and its receptor, CX3CR1, have been recently shown to be required for posttrauma cortical brain neuron microglia-mediated remodeling in a whisker lesioning paradigm (Gunner et al. 2019).

To determine if *orion* was only needed for astrocyte infiltration into axonal MB bundles or if it could have a general function in debris engulfment and phagocytosis, we examined elimination of synaptic debris by astrocytes in the VNC taking advantage of the fact that astrocytes are already present in the VNC at larval stages (Stork et al. 2014) before neuronal remodeling. Interestingly, we also show that *orion* is involved in the overall transformation of astrocytes into phagocytes. In addition, analysis of different neuronal remodeling paradigms including vCrz^+^ and PDF-Tri neurons demonstrate the role of *orion* in the elimination of structures other than MBs via ensheathing and cortex glia. Therefore, our results indicate that neuron secreted Orion activates phagocytosis in three different types of glial cells (astrocytes, cortex and ensheathing glia) and point Orion as an activator not only of glial infiltration but also of engulfment and phagocytosis.

## Material and Methods

### Drosophila stocks

All crosses were performed using standard culture medium at 25 °C. Except where otherwise stated, alleles have been described (http://flystocks.bio.indiana.edu). *Crz-GAL4* and *UAS-Casor* were provided by Jae H. Park, *alrm-GAL4* and *alrm-GAL4 UAS-GFP repoflp^6^; FRT2A, Tub-GAL80* were from Marc Freeman. We used ten *GAL4* lines: *201Y-GAL4* expressed in γ MB neurons, *PDF-GAL4* expressed in all PDF neurons, *Crz-GAL4* expressed in central and ventral Crz neurons, *Or85e-GAL4* expressed in ORNs, the pan-neuronal driver *elav-GAL4, dpr1-GAL4* (Nakamura et al. 2002), expressed in the wing nerve of wing vein L1, *NP2222-GAL4* expressed essentially in cortex glia, *MZ0709-GAL4* expressed in ensheathing glia, *alrm-GAL4* expressed exclusively in glial astrocytes (Doherty et al. 2009; Stork et al. 2014) and the pan-glial driver *repo-GAL4* expressed in all glia (Sepp et al. 2000).

### Adult brain dissection, immunostaining, and astrocyte clone visualization

Adult fly heads and thoraxes were fixed for 1 h in 3.7% formaldehyde in phosphate-buffered saline (PBS) and brains were dissected in PBS. For larval and pupal brains, brains were first dissected in PBS and then fixed for 15 min in 3.7% formaldehyde in PBS. They were then treated for immunostaining as described (Boulanger et al. 2021). Antibodies, obtained from the Developmental Studies Hybridoma Bank, were used at the following dilutions: mouse monoclonal anti-Fas2 (1D4) 1:10; mouse monoclonal anti-brunchpilot (nc82) 1:25, mouse monoclonal anti-PDF (PDF C7) 1:50, mouse monoclonal anti-discs large (4F3) 1:100 and mouse monoclonal anti-Repo (8D1.2) 1:10. Anti-Crz rabbit antibody (from Jan Veenstra) was used at 1:2000. Goat secondary antibodies conjugated to Cy3, Alexa 488, and Cy5 against mouse or rabbit IgG (Jackson Immunoresearch Laboratory) were used at 1:300 for detection, Cy3 conjugated goat anti-HRP was used at 1:100, (Jackson Immunoresearch laboratory). DAPI (Sigma) was used at 1:1000 from a 10 mg/ml stock solution. To visualize astrocyte clones in the VNC, first instar larvae were heat-shocked at 37 °C for 1 h. Adult brains were fixed for 15 min in 3.7% formaldehyde in PBS before dissection and GFP visualization. Por lysotracker staining, VNC were dissected in 1X PBS without fixation and incubated in a lysotracker solution (Lysotracher Red DND-99, Invitrogen) 1:5000 during 15 min, then washed and fixed as previously.

### Microscopy and image processing

Images were acquired at room temperature using a Zeiss LSM 780 and a Leica SP8 laser scanning confocal microscope (MRI Platform, Institute of Human Genetics, Montpellier, France) equipped with a ×40 PLAN apochromatic 1.3 oil-immersion differential interference contrast objective lens. The immersion oil used was Immersol 518F. The acquisition software used was Zen 2011 (black edition). Contrast and relative intensities of the green (GFP), of the red (Cy3), and of the blue (Cy5 and Alexa 647) channels were processed with the ImageJ and Fiji software. Settings were optimized for detection without saturating the signal. For each set of figure settings were constants. We used the Imaris (Bitplane) software to generate 3D structures of glia surrounded Crz neuron cell bodies from regular confocal images in order to determine their engulfment degree (engulfed, in contact, no contact). The images were taken using the same confocal setting within the same set of the experiments, and the data were processed in parallel. The experiments were repeated for at least two times.

### Quantitation of immunolabelling

To determine Crz cells bodies engulfment degree, we established three categories of phenotypes: “no contact,” when glial cell did not contact cell body surfaces, “in contact,” when glial cells (GFP^+^) contacted 50 % or less of the vCrz^+^ cell body surface, and “engulfed,” when >50% of vCrz^+^ cell bodies were surrounded by glial membranes. For PDF-Tri neuron surface measures were performed using the ImageJ software. The outline of the PDF-Tri staining was drawn using the freehand selection tool, on a z-stack of all the sections comprising PDF-Tri neurons in which we merged GFP and anti-PDF staining. The number of somas at each timepoint and in each condition was calculated based on confocal images. To quantify phagocytic vesicles, we counted the total amount of vesicles observed in a VNC randomly chosen confocal plan. To quantify Crz neurites we analyzed three types of GFP^+^ neurites: horizontal axons (8 tracks), medial and lateral axons (2 tracks). We considered axon was present when the GFP staining was not disrupted along the whole track. Only horizontal axons were quantified with the anti-Crz antibody, because they are shorter and easier to follow all along the process. To quantify Crz cell bodies (16 cell bodies), we considered only dots bigger than surrounding debris and located at the right emplacement on the VNC. To quantify the Casor probe GPF localization, we considered three categories: nuclei, when the fluorescence was only nuclear, cytoplasm, when only cytoplasmic GFP expression was observed and nuclei + cytoplasm when GFP expression was homogeneously distributed between nuclei and cytoplasm.

### Injury (axotomy) assay in the wing

Animals were kept at 20 °C for 5-7 days prior axotomy, unless stated otherwise. Axotomy was performed as described (Paglione et al. 2020; Llobet Rosell et al. 2022). One wing per anesthetized fly was cut approximately in the middle. Flies were returned to individual vials. 7 days post axotomy (7 dpa), wings were mounted onto a slide, and imaged with a spinning disk microscope to assess and quantify for intact or degenerated axons, as well as the remaining uninjured control axons.

### Statistics

Comparison between two groups expressing a qualitative variable was analyzed for statistical significance using the two-sided Fisher’s exact test. Comparison of two groups expressing a quantitative variable was analyzed using the two-sided nonparametric Mann–Whitney *U* test. (BiostaTGV: http://biostatgv.sentiweb.fr/?module=tests). Values of *p* <0.05 were considered to be significant. Graphs were performed using the GraphPad Prism software (version 8.1.1). Statistical significance was defined as ****p<0.0001, ***p<0.001, **p<0.01, *p<0.05, ns, for not significant. The number of neurons (n) in each group is included in a parenthesis.

## Results

### Orion is required for debris engulfment and phagocytosis by astrocytes in the pupal neuropil

In a recent work we established that *orion* was required for astrocyte infiltration into the MB g axon bundle during g axon pruning (Boulanger et al. 2021). Consequently, the phagocytosis steps succeeding infiltration such as debris engulfment, phagosome formation and debris digestion into astrocytes do not occur in the absence of Orion. Since astrocytes send extensions and already infiltrate the whole neuropil during larval stages before remodeling at pupal stages, we considered the VNC a suitable emplacement to determine if *orion* was involved in debris engulfment and phagosome formation. For this purpose, we first examine the overall morphology of the VNC located astrocytes in wild type and *orion* mutants at L3 stage using *UAS-CD8-GFP* driven by the astrocyte specific *Alrm-GAL4* (n ≥ 20) (Fig. 1A, B). In both cases, astrocytes displayed similar morphologies and locations with thin extensions infiltrating the internal side of the neuropil.

**Figure 1.**
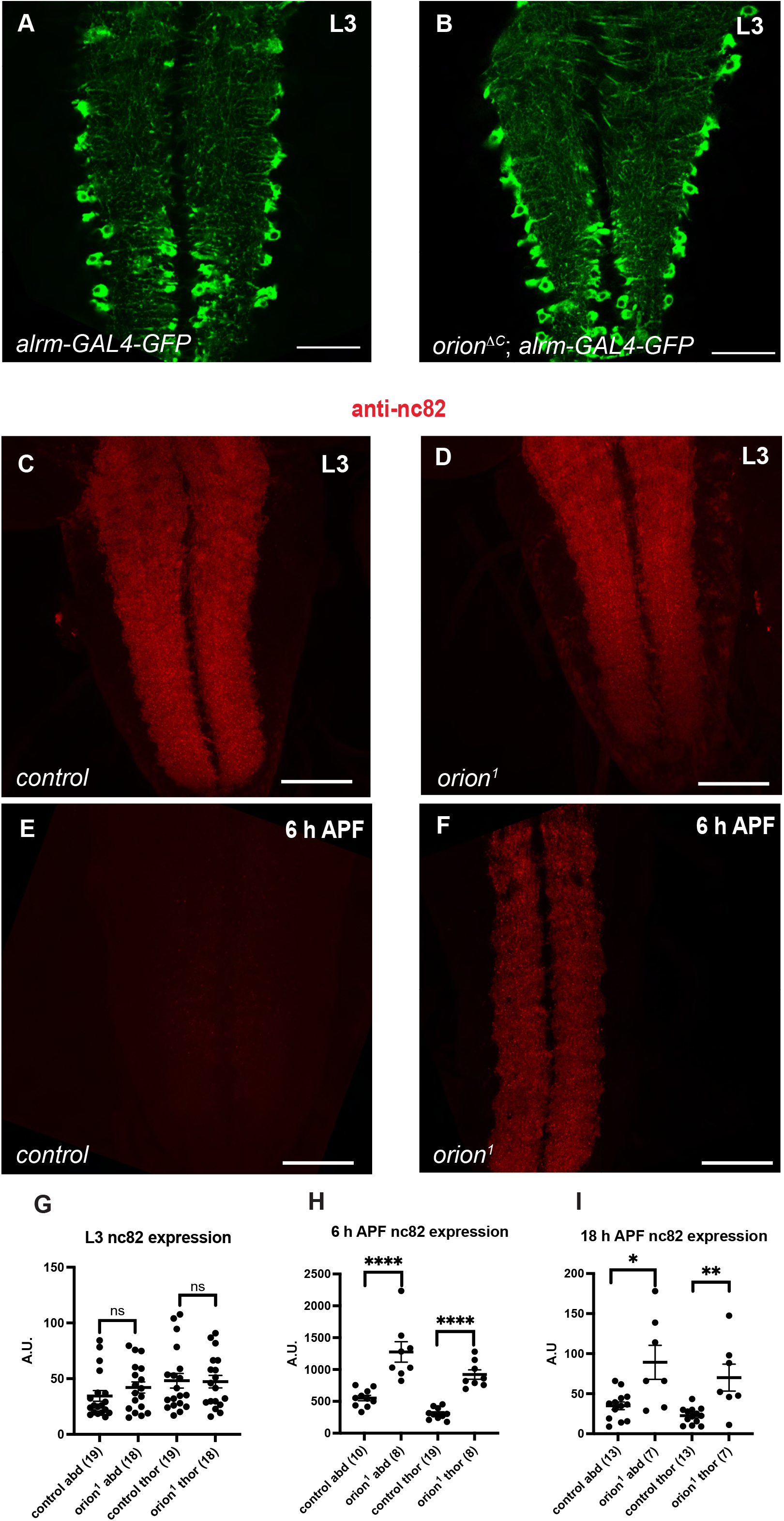
Orion is required for the elimination of synaptic material in the astrocyte-infiltrated VNC. **(A, B)** Astrocytes in controls **(A)** and *orion^ΔC^* **(B)** are visualized by the expression of *alrm-GAL4* driven *UAS-mCD8-GFP* (green) at larval stage (n ≥ 20 for each condition). Note that astrocyte extensions infiltrate the larval neuropil in both conditions. **(C-F)** Active zones were labeled with the nc82 antibody (antibody for Brp, red) at larval **(C, D)** and 6 h APF **(E, F)** stages in control **(C, E)** and *orion^1^* mutants **(E, F)** (n ≥ 20 for each condition). Note the higher amount of nc82 staining observed in F compared to wild type in E, at 6 h APF. Confocal images are z-projections. Scale bars are 40 μm in A, B and 50 μm in C-F. **(G-I)** Quantifications of expression levels of nc82^+^ puncta in arbitrary units at L3 **(G),** 6 h APF **(H)** and 18 h APF **(I)** in thorax (thor) and abdominal (abd) regions of the VNC are shown. Genotypes are listed in Supplementary list of fly strains. Results are means ± S.E.M. (Mann-Whitney *U* test).

Neuronal remodeling during metamorphosis results in the loss of nearly all synapses in the neuropil by 48 h after pupa formation (APF), before adult-specific synapses are generated. To determine if Orion had an additional role in astrocyte engulfment and phagocytosis, we used antibodies to the presynaptic active zone marker Bruchpilot (nc82) to label synapses and examine their fate at larval and pupal stages. At L3, nc82 labeled abdominal and thoracic VNC synapses throughout the neuropil at similar intensities in control and *orion^1^* (Fig. 1C, D, G). At 6 h APF nc82 staining persisted in *orion* mutant VNC compared to controls (Fig. 1E, 1F, 1 H), as well as at 18 h APF (Fig. 1I), pointing to an additional role of *orion* in the engulfment and phagocytosis of VNC synapses by the surrounding astrocytic cells. To confirm this data, we examined the astrocyte process morphology at 6 h APF in wild type and *orion* lacking flies. In wild type, VNC astrocytes displayed vacuolar structures (Tasdemir-Yilmaz and Freeman 2014) compared to *orion^ΔC^*, in which the astrocyte structure was filamentous all over the VNC with a low number of small vesicles (Fig 2 A–C). Suppression of astrocytic transformation was also observed in individual *orion* mutant astrocyte clones labeled with *mCD8-GFP* driven by *alrm-GAL4*, in which, vesicular phagocytic structures were not apparent compared to wild type (Fig. 2 D, E). Next, we sought to determine if *orion* was involved in astrocytic synapse engulfment. We clearly observed GFP+ astrocyte extensions in wild type and *orion* mutant VNCs (n ≥ 7 VNCs), but engulfed synaptic debris labeled with nc82 were only observed in wild type conditions (Fig. 2 F, G). Finally, to determine if these vesicles corresponded to phagocytic vesicles, we labeled VNCs with the phagolysosome marker lysotracker. Acidic phagocytic activity was only observed in controls at 6 h APF, reflected by a high lysotracker staining inside of the astrocytic vesicles, which was absent in the rare small astrocytic vesicles observed in *orion* lacking VNCs (Fig. 2 H, I). These results suggested that Orion is not only required for astrocytic infiltration, but also for synapse engulfment and phagocytosis and point to Orion as an overall activator of astrocyte transformation into phagocytes.

**Figure 2.**
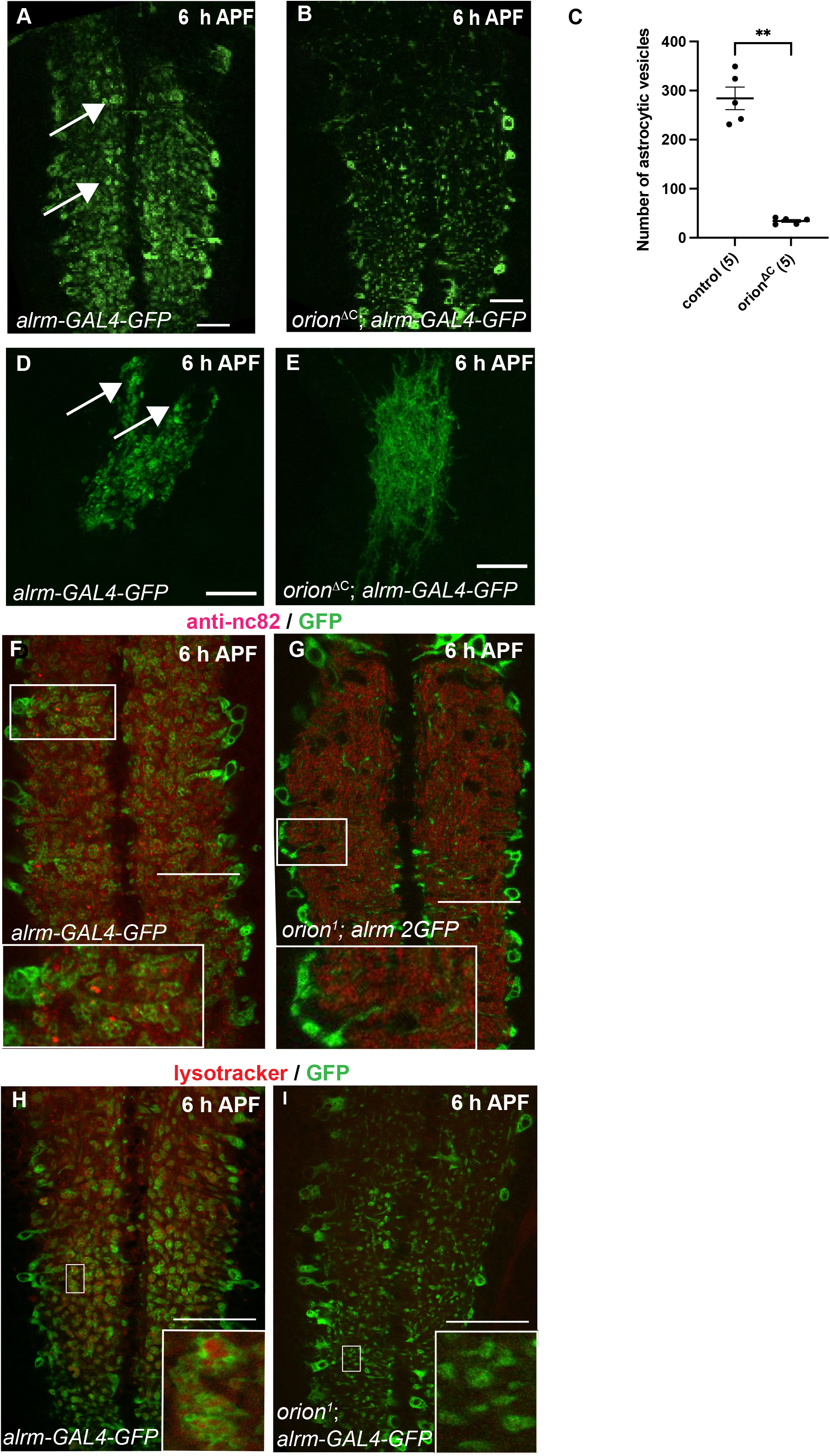
Orion regulates transformation of astrocytes into phagocytes that engulf and phagocyte synaptic material in the pupal neuropil. **(A-I)** Astrocytes are visualized by the expression of *alrm-GAL4* driven *UAS-mCD8-GFP* (green) in VNCs **(A, B; F - I)** and single astrocyte clones **(D, E)**. Red staining represents either active zones labelled with the nc82 antibody **(F, G)** or phagosomes labeled with a lysotracker staining **(H, I)**. GFP-labelled vesicular structures are pointed by arrows in **A, D**. Vesicles are mostly not observed, or very small in *orion* mutants **(B, E)**. n ≥ 20 VNC and n ≥ 10 clones in control and *orion* mutants. (**C)** Quantification of the number of astrocytic vesicles in control and *orion* mutant whole VNC. The number of VNC analyzed is included in a parenthesis for each condition. Results are means ± SEM. (**), p < 0.01 (Mann-Whitney *U* test). Insets in **F** show astrocytic vesicles containing engulfed synaptic debris absent in *orion* mutants **G**. Insets in **H** show astrocytic vesicles containing acidic phagosomes labeled with lysotracker. Note that lysotracker staining is not observed in *orion^1^* vesicles **(I)**. Confocal images are z-projections. Genotypes are listed in Supplementary list of fly strains. Scale bars are 30 μm in A, B; 20 μm in D, E and 40 μm in FI.

### Orion is required for cortex glia mediated phagocytosis of vCrz neuron cell body in the pupal neuropil

Since Orion was involved in the elimination of synaptic debris in the neuropil, we raised the question of Orion’s action in the neuropil being specific to astrocyte glia or if other types of phagocytic glial cells could also be targets of Orion. To explore this possibility, we examined the peptidergic ventral Corazonin neurons located in the ventral nerve cord (vCrz^+^), that undergo apoptosis during early metamorphosis between 0 and 6 h APF and whose cell bodies and neurites, projecting into the neuropil are eliminated (Choi et al. 2006). vCrz^+^ system is composed of 8 pairs of neurons (16 cell bodies) and three different types of neurite tracks: 8 horizontal neurites connecting cell bodies from the same segment, 2 medial vertical neurite tracks in the center of the neuropil and 2 lateral neurite tracks connecting vertically cell bodies from contiguous hemi segments. Interestingly the vCrz^+^ neurites are eliminated by astrocytic glia but their cell body seems to be removed by other types of glia, most probably cortex glia (Tasdemir-Yilmaz and Freeman 2014). Thus, we considered these neurons as a valuable model to determine if *orion* was involved in the elimination of cell bodies by glial cells other than astrocytes. In controls, at L3, the anti-Crz antibody recognizing the Crz neuropeptide labels eight pairs of vCrz neurons in the VNC (and a few neurons in the brain). By 6 h APF, almost all of the vCrz^+^ neurons are cleared from the neuropil (Choi et al. 2006). In VNCs lacking *orion (orion^1^* and *orion^ΔD^*) we found no differences in the anti-Crz staining at L3 stages compared with wild type larvae. However, most of the cell bodies and neurites persisted at 4 h APF in *orion* mutants, which was not the case in controls. Furthermore, many vCrz^+^ cell bodies and a high amount of neurite debris remained also at 6 h APF in *orion* mutants (Fig 3 A–J). These results were confirmed by driving GFP in vCrz^+^ neuronal membranes by a Crz neuron specific GAL4 (Supplemental Fig 1). We observed that neurite and cell body membrane GFP-labelled debris remaining until 18 h APF in VNCs lacking *orion* (Supplemental Fig 1 E, J). Forced expression of *orion* in neurons using the pan-neuronal *elav-GAL4* driver at 6 h APF rescued the *orion* mutant cell body and neurite phenotype (Fig. 3 K–N), indicating that expression of *orion* in neurons is sufficient for Crz neuron remodeling. In order to determine if neuronal *orion* was activating vCrz^+^ surrounding glia, leading to cell body and neurite glia-mediated phagocytosis, we analyzed the morphology of glial cells located in the vicinity of the vCrz^+^ cell bodies using the pan-glial *GAL4* driver *Repo*. A significantly higher amount of vCrz^+^ cell bodies were engulfed by glial cells in wild type compared to *orion^1^* at 6 h APF (53% over 11% respectively) (Fig. 3 P). Interestingly, in *orion* mutants, 42% of cell bodies were not in contact with glia (Fig. 3 O, P). These results suggested that vCrz^+^ neuron secreted Orion is required for the engulfment of vCrz^+^ cell bodies and neurites by glial cells. We next sought to determine which type of phagocytic glia was involved in the elimination of vCrz^+^ cell bodies and neurites. For this purpose, we used several *GAL4* lines: *alrm-GAL4, NP2222-GAL4* and *MZ0709-GAL4*, specific respectively for astrocytes, cortex glia and ensheathing glia and analyzed *UAS-GFP* driven expression at 4 h APF. As expected for *alrm-GAL4*, astrocyte extensions infiltrating the neuropil were covered by vesicular structures containing vCrz^+^ debris. These astrocytic extensions did not contact vCrz^+^ cell bodies (Fig 4 A–C) suggesting that astrocytes were not required for the elimination of vCrz^+^ cell bodies. We then assayed cortex glia specific staining and observed that cortex glia extensions contact and engulf vCrz^+^ cell bodies (Fig. 4 D, E). In contrast, ensheathing glia extensions were not observed neither around the vCrz+ cell bodies nor the vCrz+ neurites (Supplemental figure 2). These results point to the fact that Orion is a neuronal signal that allows communication not only between neurites and astrocytes but also between neuron cell bodies and cortex glia. This interaction is likely necessary to allow elimination of cell bodies after apoptotic cell death induction.

**Figure 3.**
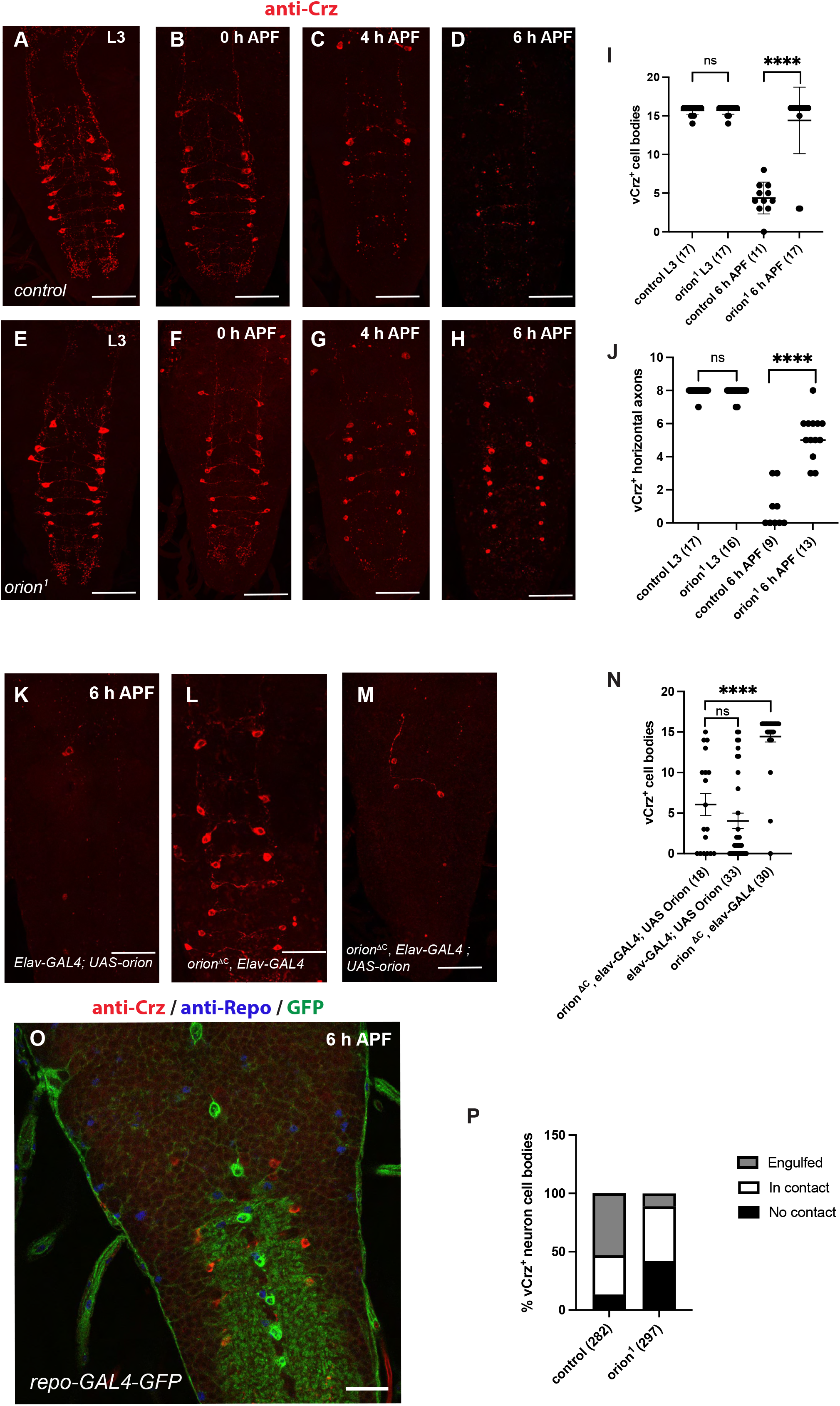
Orion is required for the elimination of vCrz^+^ cell bodies and neurites. **(A-J)** vCrz^+^ neurons were labeled with an anti-Crz antibody (red) at the indicated time points at larval (L3) and pupal stages in controls **(A-D)** and *orion^1^* mutants **(E-H)**. **(I)** Quantification of vCrz^+^ cell bodies. **(J)** Quantification of vCrz^+^ horizontal axons. **(K-M)** 6 h APF vCrz^+^ neurons were labeled with anti-Crz (red) in controls **(K)** and orion mutants **(L).** Neuronal expression of *orion* rescued the *orion^ΔC^* phenotype **(M)**. **(N)** Quantification of vCrz^+^ cell bodies. **(O)** Expression of *UAS-mCD8-GFP* under the control of *Repo-GAL4* (green), anti-Crz staining (red) and anti-Repo staining (blue) are shown in 6 h APF neurons. Genotypes are listed in Supplementary list of fly strains. Error bars represent ± SEM (Mann-Whitney *U* test). **(P)** Quantification of vCrz^+^ cell bodies engulfed, in contact and not in contact with glial extensions. n values are indicated in a parenthesis for each condition. Scale bars are 70 μm in A-H and 40 μm in K-M and O.

**Figure 4.**
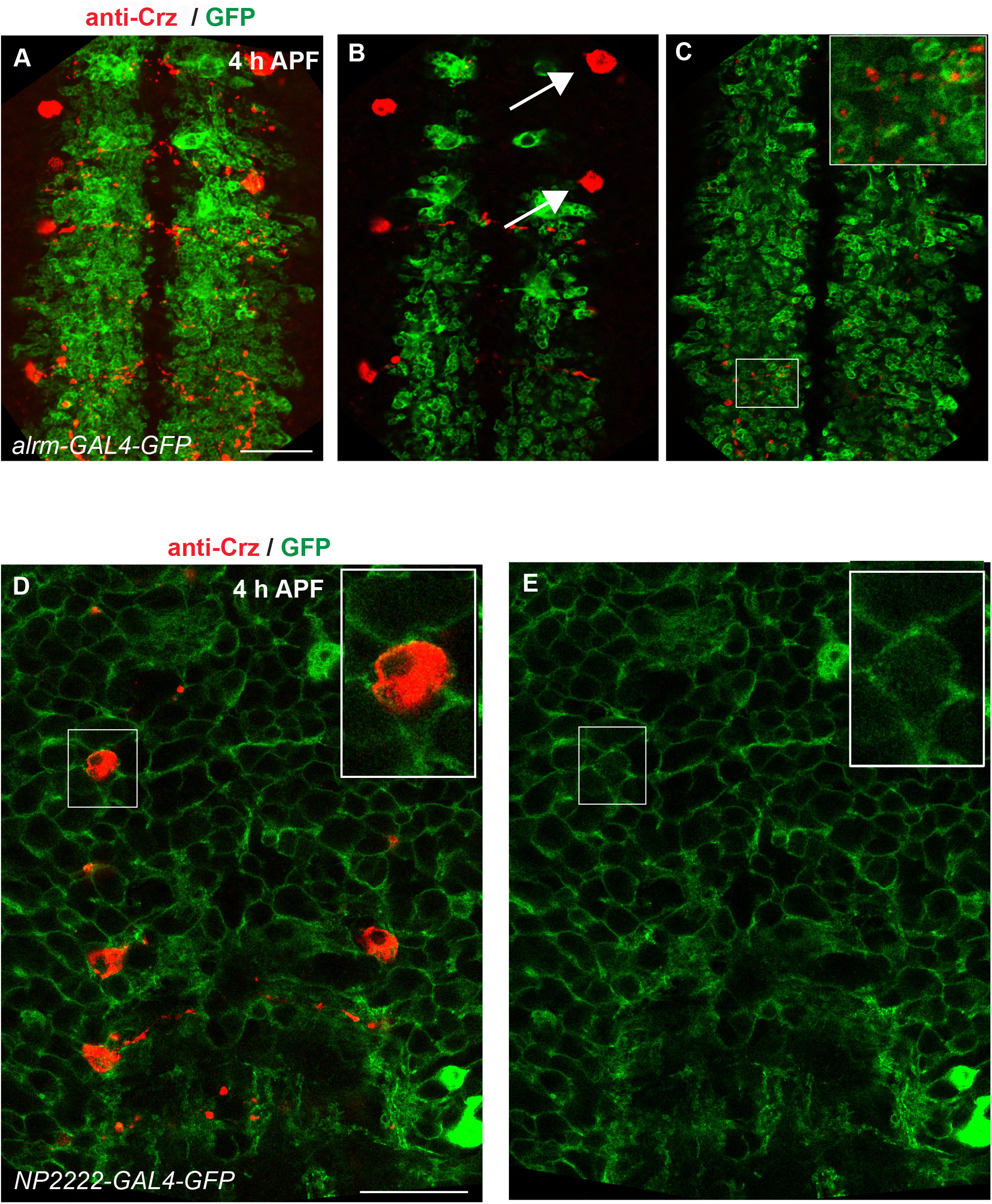
The elimination of vCrz^+^ neurites and cell bodies is mediated by astrocytes and cortex glia respectively. **(A-C)** Astrocytic glia visualized by the expression of *alrm-GAL4* driven *UAS-mCD8-GFP* (green) and vCrz^+^ neurons by an anti-Crz antibody (red) at 4 h APF. **(B)** Arrows point to vCrz^+^ cell bodies which are not reached by astrocytic extensions. **(C)** Astrocyte extensions engulf vCrz^+^ debris. See inset for higher magnification. **A** is a z-projection confocal image, **B** and **C** are single confocal plans from A. **(D, E)** Confocal plans showing cortex glia visualized by the expression of *NP2222-GAL4* driven *UAS-mCD8-GFP* (green) and vCrz^+^ neurons by an anti-Crz antibody (red) at 4 h APF. Insets illustrate a vCrz^+^ cell body being engulfed by cortex glia. Genotypes are listed in Supplementary list of fly strains. Scale bars are 30 μm.

To determine if cortex glia had an active role in the elimination of the vCrz^+^ cell bodies during vCrz^+^ apoptosis, we compared control and *orion* lacking VNCs expressing the GFP-based caspase sensor (*UAS-Casor*) probe in vCrz^+^ neurons. This probe is designed to change its subcellular localization from the cell membrane to the nucleus following proteolytic cleavage by active caspases and allows to monitor differences in caspase activity in neurons like the vCrz^+^ (Lee et al. 2018). If *orion* facilitated apoptosis after caspase activation we would anticipate a delay in the nuclear localization of the Casor probe in *orion*-lacking VNCs compared to wild type VNCs during development. However, no difference was observed between wild type and o*rion*-lacking VNCs (Supplemental figure 3), neither at 0 h APF, in which the overall vCrz^+^ soma GFP+ staining was cytoplasmic (78 % and 86 % for controls and *orion* mutants respectively), nor at 2 h APF, where the cords displayed a mostly nuclear GFP+ staining (90 % and 100 % for controls and *orion* mutants respectively). These data suggested that Orion does not act on the vCrz^+^ neuron cell autonomous caspase activity leading to vCrz^+^ neuronal cell body apoptosis.

### Orion is required for elimination of PDF-Tri neurons by ensheathing and cortex glia in new born flies

We used an additional paradigm in order to study ensheathing glia phagocytic potential effects during remodeling, the peptide-dispersing factor tritocerebrum (PDF-Tri) neurons, which are characterized as a developmentally transient population that initially appears during mid-pupal development and undergoes programmed cell death soon after adult eclosion (Helfrich-Forster 1997; Gatto and Broadie 2011). PDF-Tri neurons consist of 1–2 pairs of somata bilaterally positioned in the tritocerebrum on the anterior face of the brain, in close proximity of the esophagus (Vita et al. 2021). PDF-Tri processes extend posteriorly and ventrally into the suboesophageal zone (SEZ), dorsally through the distal medial bundle (MDBL), and into the superior medial protocerebrum near the dorsal brain surface (Fig. 5A). The highly ramified dendritic arbors and axonal projections of the PDF-Tri neurons project throughout the SEZ just dorsal to, and surrounding, the esophageal foramen (Fig. 5A). As PDF-Tri processes project toward the posterior surface through the MBDL, two linearized branching tracks extend laterally toward the dorsal side of the brain (Fig. 5A). Recent studies have shown that these neurons fail to be removed by blocking glia phagocytosis (Vita et al. 2021). In addition, it was reported that cortex and ensheathing glia are the major phagocytic glial cells responsible for the elimination of their cell bodies and axonal processes respectively. Consequently, the PDF-Tri neurons represent a suitable structure to establish if Orion has a role on their cell body and neurite elimination *via* the surrounding cortex and ensheathing glia respectively. To examine PDF-Tri neurons, neurons were co-labeled with *PDF*-*GAL4*-driven membrane-tethered green fluorescent protein (GFP) and an anti-PDF (Fig 5); which label not only the PDF-Tri neurons but also a network of PDF surrounding positive neurons (Supplemental Fig 4 A, B). In newly eclosed adults (0 days), PDF-Tri neurons are consistently present along the brain central midline in controls and *orion^1^* mutants. Dense PDF+ projections occur in the SEZ and surrounding the esophageal foramen. Distribution of fluorescence and PDF-Tri staining overall were also similar in control and *orion^1^* mutants. In addition, a similar amount of cell bodies was observed (Figure 5, A–D). Three days later (3 days post-eclosion), extensive removal of the PDF-Tri circuitry is underway, with the disassembly and loss of axons, dendrites and cell bodies in control animals as reported previously (Gatto and Broadie 2011). However, the neurite circuitry and cell bodies are maintained in *orion^1^* (Figure 5, E–H). Surrounding PDF+ neuronal circuitry which is not fated to remodel is not affected neither in control nor in mutant animals (Supplemental Fig 4 A, B). Both the *PDF*-*GAL4*-driven GFP and anti-PDF labeling of the PDF-Tri neurons disappear with a similar time course, indicating elimination of both the developmentally transient PDF-Tri neuronal membranes and the cytoplasmic PDF peptide content at this stage. By one week post eclosion (Fig 5, I–K), PDF-Tri cell bodies and most of the axonal and dendritic processes were still present in *orion^1^* mutants compared to controls, and they occupied mostly the initial surface. Finally, we observed that PDF-Tri neurites persisted in *orion* lacking flies at three weeks after eclosion (Supplemental Fig 4 C-E).

**Fig 5.**
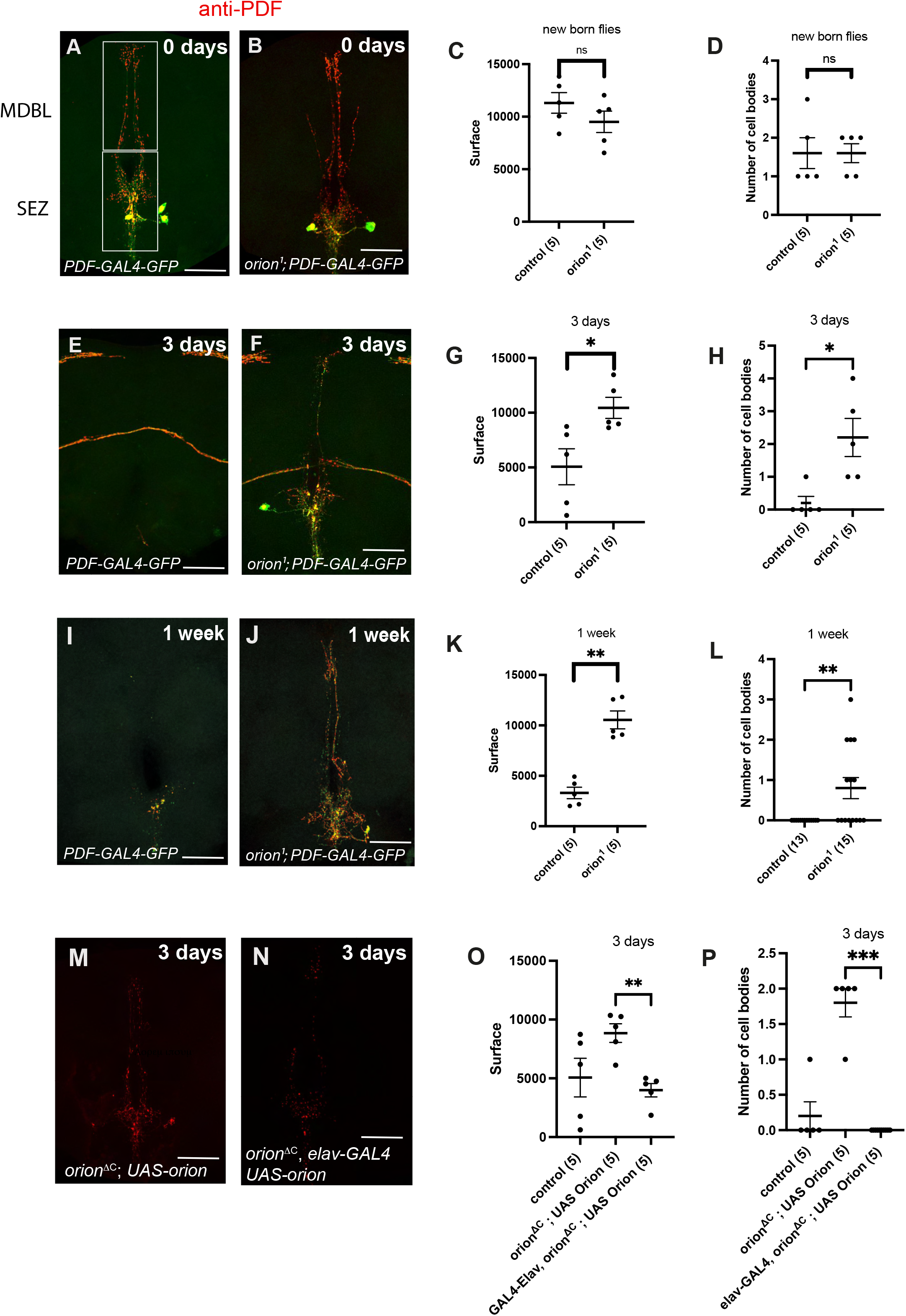
Orion mutants retain developmentally transient PDF-Tri neurons. **(A, B; E, F; I, J)** Confocal z-stacks showing PDF-Tri neurons visualized by the expression of *PDF-GAL4* driven *UAS-mCD8-GFP* (green) and labeled with anti-PDF antibody (red) at the indicated time points in controls **(A, E, I)** and *orion^1^* mutants **(B, F, J).** Genotypes are listed in Supplementary list of fly strains. Scale bar is 50 μm. **(C, G, K)** Surface (in μm^2^) occupied by the PDF-Tri arborization at 0 days (new born flies), 3 days and one week respectively in controls and *orion* mutants. **(D, H, L)** PDF-Tri cell bodies at 0 day, 3 days and one week in controls and *orion* mutants. **(M, N)** Confocal z-stacks showing PDF-Tri neurons labeled with an anti-PDF antibody at 3 days after birth in *orion^㥌^* mutants **(M)** and in *orion^ΔC^* rescued by the expression of *elav-GAL4* driven *UAS-orion* **(N). (O, P)** Rescue quantifications of PDF-Tri surface (in μm^2^) (O) and number of cell bodies (P) are shown. Genotypes are listed in Supplementary list of fly strains. Scale bar is 50 μm. Error bars represent ± SEM (Mann-Whitney *U* test). n values are indicated in a parenthesis for each condition.

Forced expression of *orion* in neurons using the *elav-GAL4* driver three days after birth rescued the *orion* mutant cell body and neurite phenotypes (Fig. 6 M–P), suggesting that expression of *orion* in neurons is sufficient for PDF-Tri neuronal remodeling.

**Fig 6.**
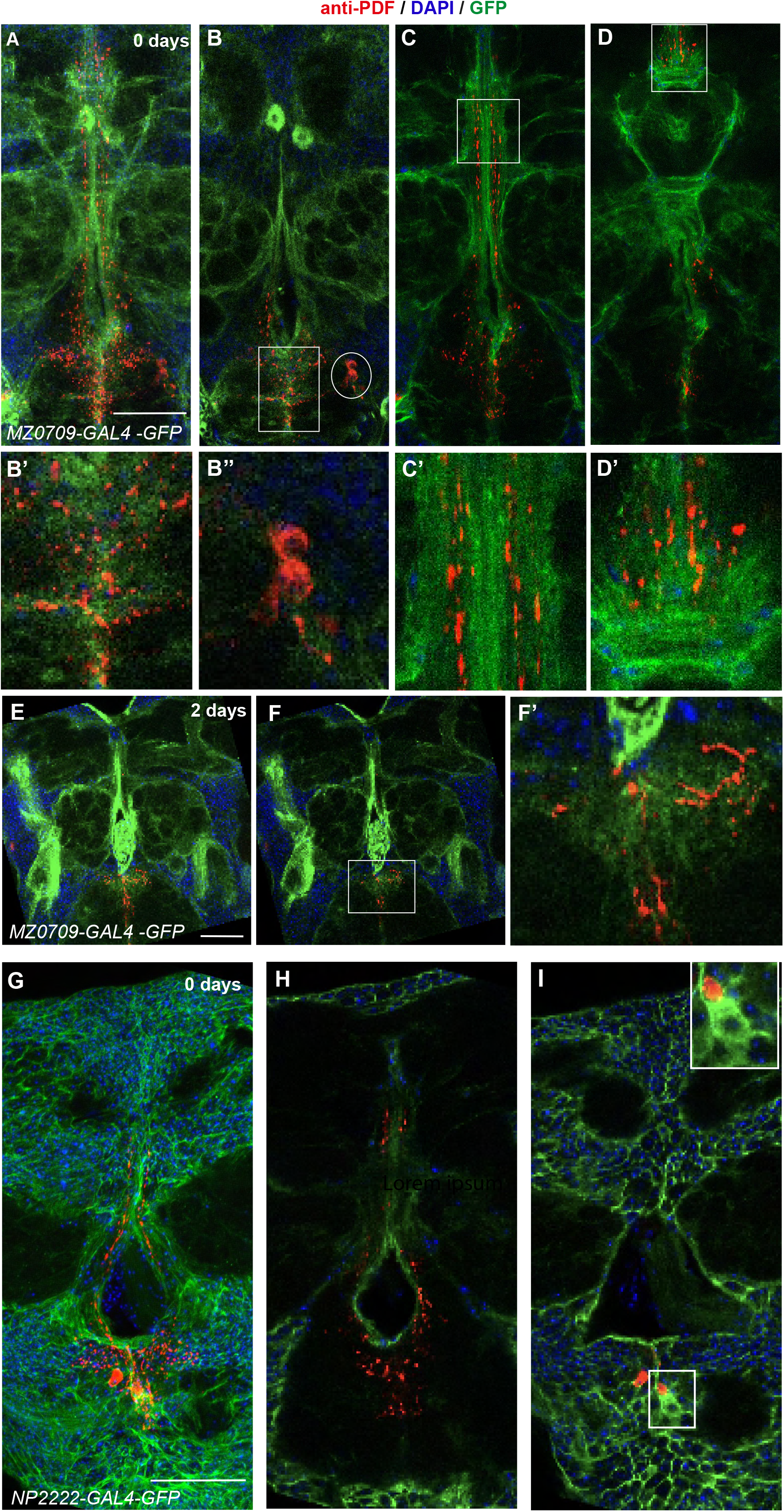
Orion mediates PDF-Tri neuron elimination *via* cortex and ensheathing glia. **(A, E)** Confocal z-stack showing ensheathing glia visualized by the expression of *MZ0709-GAL4* driven *UAS-mCD8-GFP* (green) and labeled with anti-PDF antibody (red) and DAPI (bleu) at 0 **(A)** and two **(E)** days. Different regions of the brain contained in these z-stacks are shown as single confocal plans (B-D for z-stack A, F for z-stack E). SEZ region is included in a rectangle in **B** and **F**, MBDL region is included in a rectangle in **C** and top of the MBDL region is included in a rectangle in **D**. These regions are shown at higher magnification in **B’, F’, C’ and D’** respectively, showing PDF-Tri neurites engulfed by ensheathing glia. Round in **B** encircles a PDF-Tri cell body devoided of ensheathing glia, which does not engulf PDF-Tri cell bodies (higher magnification is in B’’). **(G)** Confocal z-stack showing cortex glia visualized by the expression of *NP2222-GAL4* driven *UAS-mCD8-GFP* (green) and labeled with anti-PDF antibody (red) and DAPI (bleu) at 0 days. **(H, I)** Single confocal plans showing PDF-Tri dendrites **(H)** and cell bodies **(I)** contained in the G image stack. Note the absence of PDF-Tri dendrite surroundered by cortex glia in H and the high amount of cortex glia extensions present around the PDF-Tri cell bodies in I. **(I)** A cortex glia cell phagocyting a PDF-Tri cell body (red) is included in the region enclosed by the square. (n ≥ 5 for each condition). Genotypes are listed in Supplementary list of fly strains. Scale bars are 50 μm.

To bolster whether Orion signals to glia to initiate phagocytosis for PDF-Tri neuron elimination, both wild type and *orion^1^* animals were imaged using specific glial markers and the anti-PDF antibody at 0 and 2 days post-eclosion, when cell bodies and neurites still are respectively present. Cortex and ensheathing glia are both primary phagocytes of PDF-Tri neurons (Vita et al. 2021). Concerning astrocytic glia, its role was not evidenced in PDF-Tri phagocytosis, but since Orion was shown to essentially signals to astrocytes in the MBs and the VNC we also included them in our analysis. Selective drivers were used for the astrocyte-like (*alrm-GAL4*), ensheathing (*MZ0709*-*GAL4*), and cortex (*CG2222*-*GAL4*) glia. Combining *mCD8-GFP* under control of each driver generates distinctive brain localization and cellular morphology for each glial class in regions surrounding PDF-Tri neurons. GFP-labeled astrocytes exhibited a small cytoplasm size and some long and ramified extensions (Supplemental fig 5). Astrocytes did not surround somas (Supplemental fig 5 A, B arrow) or processes (Suppl Fig 5 A, C, D) and their cytoplasm did not show phagocytic vesicles. In addition, no engulfment of debris neither at 0 day nor at 2 days was observed. Astrocytic extensions become longer at 2 days, but, they did not seem to phagocyte PDF-Tri debris (Supplemental Fig 5 E, F).

Next, we looked at ensheathing glia GFP+ driven membranes that prominently define and surround each brain neuropil at 0 and 2 days (Fig 6 A–F’). PDF-Tri dendrites are surrounded by GFP+ ensheathing glial membranes at 0 day (birth) (Fig. 6, A, B) as well as both of the PDF-Tri vertical tracks (Fig 6 A, C) and the distal PDF-Tri neurites (Fig 6 A, D). Ensheathing glia extensions are not observed around PDF-Tri cell bodies (Fig 6 B). Similar results were observed in 2 days brains (Fig 6 E–F’).

Concerning cortex glia, we observed cortex glia extensions phagocyting individual PDF-Tri cell bodies (Fig 6 I), but not neurites (Fig 6 H) at 0 day after eclosion, suggesting that Orion mediates PDF-Tri cell body phagocytosis.

As well as the vCrz^+^ cell bodies, we did not observe any active role of glia in the PDF-Tri cell body apoptosis (Supplemental Fig 6). These data suggest that Orion signals to both ensheathing and cortex glia allowing PDF-Tri neuronal debris phagocytosis, with the cortex glia class acting specifically on cell body clearance while the ensheathing glia class facilitates debris elimination more distally.

### Orion is dispensable for the debris clearance of severed ORN axons by ensheathing glia after maxillary palp ablation

A well-established system to study communication between neurons remodeling and ensheathing glia is based on the analysis of ORNs and surrounding ensheathing glia after the ablation of maxillary palps or antennae (Doherty et al. 2009). ORN cell bodies are housed in the third antennal segment and maxillary palps of adult *Drosophila* with axons projecting to the antennal lobe of the brain via the antennal and maxillary nerve, respectively.

After axotomy, the severed axon separated from its neuronal cell body degenerates by an evolutionarily conserved axon death signaling cascade, and the resulting debris is cleared by surrounding glia, commonly known as Wallerian degeneration (Llobet Rosell and Neukomm 2019). The surgical ablation of the antennae or maxillary palps severs ORN axons that project into the brain, triggering Wallerian degeneration (Doherty et al. 2009). Ensheathing glia infiltrate the injury site and engulf ORN axonal debris. In control animals, GFP^+^ axonal debris was cleared from the brain, while virtually all GFP^+^ degenerating ORN axons persisted in several mutants in which glial phagocytosis was blocked (Doherty et al. 2009; Musashe et al. 2016). Thus, this is a model of interest to analyze lack of *orion* effects. We used flies expressing membrane-tethered GFP in a subset of maxillary palp ORNs (*Or85e-GAL4*). Our qualitative ORN structural analysis showed that in uninjured animals (control), ORN axons (labeled with mCD8-GFP) had a smooth morphology as they projected across the antennal lobe, and GFP intensity in glomeruli was very strong and presented membrane continuity in all the brains analyzed. These observations were similar in wild type and *orion^1^* (Supplemental Fig 7 A, B). At 24 h after palp ablation we observed discontinuity in axon fibers (dotted staining) outside of the glomeruli reflecting a high level of fragmentation induced Wallerian degeneration in both control and *orion^1^* (Supplemental Fig 7, C and D) and only traces of GFP^+^ axonal debris from glomeruli and fibers were present in wild type and *orion^1^* mutant brains three days after palp ablation (Supplemental Fig 7 F, G). These results suggested that Orion is not required for the elimination of ORNs neurons by ensheathing glia after maxillary palp injury.

### Orion is dispensable for nerve debris removal by wrapping glia following wing injury and NMJ dismantling

The wing serves as another well-established model to study the communication between severed axons and surrounding glia. The *Drosophila* marginal L1 wing vein allows for a graded level of axotomy, which results in two populations of axons: severed axons, whose cell bodies are distal to the injury site; and intact axons, whose cell bodies are proximal to the injury site (Neukomm et al. 2014). The axonal debris of severed mechanosensory and sensory neurons is cleared by wrapping glia (Neukomm et al. 2014). Sensory neuron MARCM clones in the wing expressing membrane-tethered GFP *(UAS-mCD8-GFP)* driven by *dpr1-Gal4* were subjected to axotomy, in wild type and *orion^1^* heterozygous females, as well as wild type and *orion^1^* hemizygous males. 7 days after axotomy (7 dpa), in the injured wing, the uninjured (control) axons, axonal debris traces, and severed intact axons were scored. Age-matched uninjured wings served as controls.

We found no extra severed intact axons at 7 dpa, suggesting that the execution of the axon death program is not affected by the *orion^1^* mutation (Figure 7). We also found that the resulting axonal debris was cleared to a similar extent in all genotypes. It is in stark contrast to the *draper^Δ5^* mutation that impairs glial clearance and results in a full penetrance of persistent axonal debris (Neukomm et al. 2014). It is worth mentioning that the MARCM clone numbers were higher in males compared to females. However, clone numbers do not have any impact on the outcome of the injury assay. Together, these observations suggest that *orion* is dispensable for the wrapping glia-mediated clearance of axonal debris in the L1 wing vein.

**Fig 7.**
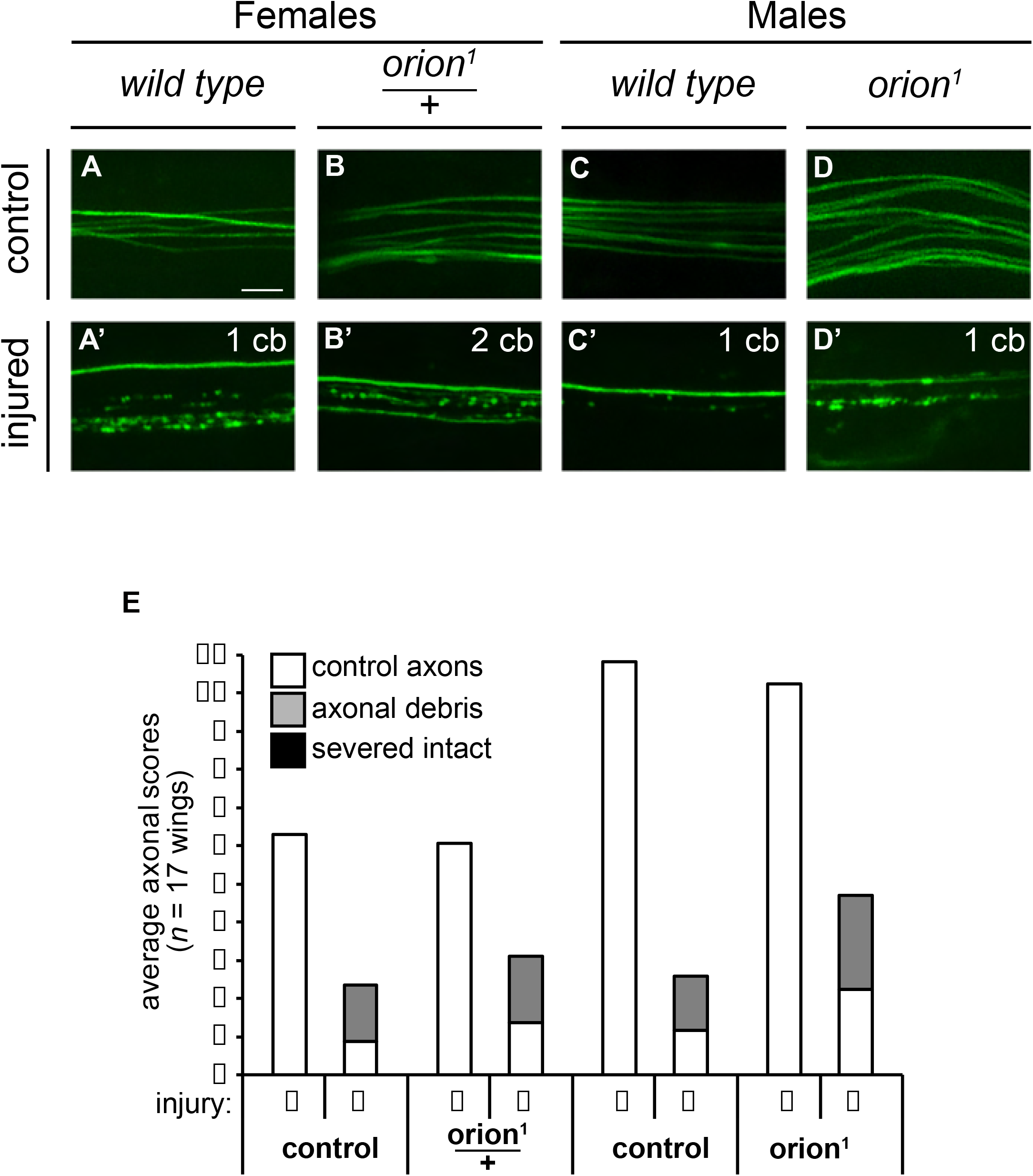
Orion is dispensable for clearing axonal debris after axotomy in the wing. **(A-D’)** Images are confocal z-stacks showing L1 vein axons labelled by *dpr1-GAL4* driven *UAS-mCD8-GFP* (green). Representative pictures of control and injured (7 dpa) axons in wild type, heterozygous *orion^1^*, and hemizygous *orion^1^* animals are shown. (E) Quantification of average axonal scores of uninjured controls, debris, and severed intact axons (white, gray, and black, respectively). MARCM clones were generated in animals of the indicated genetic background. Scale bar is 5 μm.

In further analysis we explored the effect of Orion on wrapping glia during neuromuscular junction (NMJ) developmental dismantling. NMJ dismantling during early stages of metamorphosis was previously described as a process involving retraction of wrapping glia preceding motor neuron retraction (Boulanger et al. 2012). Even though phagocytic wrapping glia was not observed during this process, it was shown that disrupting glial function blocks NMJ dismantling. Suggesting that when the glial dynamics is blocked NMJ dismantling may be also blocked. Consequently, we investigated a potential role of motor neuron secreted Orion that could affect NMJ dismantling by allowing wrapping glia dynamics. To do this, we studied muscle 4, at abdominal segment 3, NMJ development in larva and at 6 h APF. Presynaptic motor neuron membranes were labeled with anti-HRP and postsynaptic muscle with anti-discs-large (DLG). Anti-HRP and anti-DLG staining of larval and pupal NMJs showed well defined and organized synaptic boutons at L3 in control and orion^1^ mutants in 100% of the analyzed NMJ (n = 31 and 38 for wild type and *orion^1^* mutants respectively) (Supplemental figure 8 A-F). By 6 h APF the NMJs appeared completely disorganized in both, wild type and *orion* lacking animals (Supplemental figure 8 G-L), thus preventing the distinction of individual synaptic boutons. Only the presence of retraction bulbs was observed, pointing to normal synaptic retraction in both control and *orion^1^* mutants. In addition, the postsynaptic components labeled with the anti-DLG antibody became fuzzy and the DLG staining appeared completely fragmented and often absent in 90.3% of wild type NMJ versus 89.2% of orion^1^ NMJ (n = 31 and 38 wild type and orion^1^ mutants respectively). Thus, the similar NMJ morphology observed in control and *orion^1^* mutants, suggested no effect of Orion in NMJ surrounding wrapping glia.

## Discussion

### Role of *orion* in engulfment and phagocytosis by astrocytes

Developmental remodeling of neural circuitry is a key strategy employed to prune redundant, inappropriate, or interfering neurons in order to optimize connectivity (Wilton et al. 2019). Signals sent from dying neurons or neurites to be removed are received by appropriate glial cells. After receiving these signals, glia infiltrate degenerating sites, engulf and clear neuronal debris through phagocytic mechanisms. There are few identified signals involved in neuron-glia communication, which induce the transformation of glial cells into phagocytes during neuronal remodeling in *Drosophila* (Boulanger and Dura 2022). Many of these signaling pathways are conserved in mammals. We recently identified the chemokine-like Orion as a ligand, secreted by *Drosophila* MB g neurons of the central brain, inducing astrocytic infiltration into the γ axon-bundle (Boulanger et al. 2021). Thus, in *orion*-lacking flies, astrocytes are unable to infiltrate the γ axon bundle and consequently axon fragmentation resulting debris are not eliminated during metamorphosis. Previous observations have shown a high elimination of neuronal processes and synaptic terminals along the VNC early in development by astrocytes (Tasdemir-Yilmaz and Freeman 2014) which already infiltrate the neuropil before neuronal remodeling (Stork et al. 2014), differing from what occurs in MB bundles. In this study we show that *orion* has a key role in engulfment and phagocytosis of VNC neuronal debris. These data rise the possibility that Orion not only orchestrates glia infiltration into axonal bundle fated to degenerate but it is also required for engulfment of remnant debris and phagocytosis. One possibility is that Orion activates a pathway that allows glial membrane recruitment leading to the formation of glial extensions and the phagocytic cap (Ziegenfuss et al. 2012; Lu et al. 2014). In addition, neuron-secreted Orion might also be required for the different steps of phagosome maturation or for phagosome-lysosome association leading to the formation of phagolysosomes in astrocytes. In support of this model, proteomic analysis of the *Drosophila* phagosome from cultured *Drosophila* cells identified Orion along with known proteins involved in phagosome maturation (Stuart et al. 2007).

### *orion* mediates the transformation of astrocytes into phagocytes

Little is known about the pathways involved in glia activation. Astrocyte activation into phagocytes have been previously documented (Morizawa et al. 2017; Lee and Chung 2021; Konishi et al. 2022). Glia activation is characterized by the presence of enlarged processes and abundant phagocytic vacuoles displaying phagolysosomal activity in the astrocyte cytoplasm. This activation depends on the steroid hormone 20-hydroxyecdysone (ecdysone) receptor expression *(EcR)*, which in turn regulates the expression of *drpr*. Thus, loss of EcR signaling is sufficient to cell-autonomously suppress the transformation of astrocytes into phagocytes at pupariation (Tasdemir-Yilmaz and Freeman 2014). Interestingly, we observed a similar phenotype of astrocyte vacuolated appearance and thick extensions in *orion*-lacking flies, suggesting that Orion mediates the overall transformation of astrocytes into phagocytes leading to axon-bundle infiltration (when required), engulfment and phagocytosis.

### *orion* mediates the elimination of peptidergic neurons by cortex and ensheathing glia

MB g neurons prune their medial and dorsal axon branches and dendrites at early pupal stage, while their cell bodies remain. Later, at mid-pupal stages, they re-extend medial axon branches to establish adult-specific connectivity (Lee et al. 2000; Watts et al. 2004). In contrast, peptidergic vCrz^+^ and PDF-Tri neurons exhibit complete neurite degeneration, and their cell bodies undergo apoptotic death and are eventually completely eliminated. This elimination occurs at different developmental stages: the vCrz^+^ neuron apoptosis initiates in early pupae, whereas the PDF-Tri neuron apoptosis starts early after adult eclosion (Helfrich-Forster 1997; Choi et al. 2006; Gatto and Broadie 2011; Selcho et al. 2018). We show here that Orion is involved in the elimination of both types of peptidergic neurons, extending the role of *orion* to the elimination of apoptotic neurons. Interestingly, *orion* is needed not only for the elimination of vCrz^+^ and PDF-Tri neurites but also cell bodies. This suggest that Orion can also be presented to phagocytic cells by soma to be removed. Furthermore, since PDF-Tri neurons are eliminated in newly eclosed flies, our results formally extend the role of *orion* to young adult stages. We also observed in this study that *orion* does not seem to be involved in the cell autonomous neuronal apoptosis of these neurons as vCrz and PDF-Tri neuron apoptosis, monitored with the caspase sensor Casor, is similar in *orion* mutant and wild type flies. This suggests that, as previously reported for MB, *orion* does not act on the initial neuron-intrinsic developmental degeneration or apoptosis pathways, but rather in secondary non-cell autonomous pathways.

Cortex glia seems essential for the elimination of both types of neuronal cell bodies as revealed by the high level of cortex glia-engulfment around the vCrz and PDF-Tri cell bodies in wild types compared to *orion* mutants. Interestingly, we observed a high amount of ensheathing glia extensions along the PDF-Tri neurites, on both, the MDBL and SEZ regions. The proximity of this type of glia to PDF-Tri axons in wild type and *orion*-mutant flies, does not allow to determine the engulfment degree of these axons. Nevertheless; since the lack of Orion induces a high amount of unungulfed axonal PDF-Tri debris and based on our observations showing

PDF-Tri debris surrounded by ensheathing glia in wild type flies during PDF-Tri neuronal remodeling, we can anticipate that this type of glia is also a target of Orion. This idea is reinforced by the study of Vita *et al*. (Vita et al. 2021) suggesting that ensheathing glia drive the clearance of PDF-Tri neurons. Together these data provide evidence that Orion not only is able to signals *via* astrocyte glia but also *via* cortex and ensheathing glia depending on the neuronal remodeling paradigm.

### *orion* is dispensable in two adult axonal injury paradigms

Recent studies have shown that *orion* is required for the elimination of larval dendritic arborization (da) dendrites by phagocytic epidermal cells after laser ablation at larval stages (Ji et al. 2022). To extend the role of *orion* to axonal injury paradigms, we explored two distinct antennal and wing axotomy models in *Drosophila*. Antennal ablation triggers ORN axon degeneration and the resulting debris is eliminated by ensheathing glia (MacDonald et al. 2006; Doherty et al. 2009; Logan et al. 2012). Orion is not involved in the clearance of the ORN axonal debris, as a similar level of axon debris clearance is observed in controls and *orion^1^* mutants after ORN axotomy. In the *Drosophila* L1 wing vein, wrapping glia eliminate the debris from injured axons. After axotomy, Orion is not involved in the communication between severed axons and wrapping glia, suggesting that *orion* is not required for proper glial clearance after axotomy in adult flies, which contrasts the dendrite injury model in larvae. Thus, *orion* plays a crucial role during development, but not in adult flies (see Table I). In addition, differences in glial subtypes executing phagocytic function during development versus the adult could result from modification of glial genetic programs during development *versus* mature glia or indicate key differences in the molecular nature of remodeling or larval neurites compared with mature adult neurites.

**Table 1.** Neuronal paradigms in which the *orion* interaction with glial cells has been investigated during development and injury. Parenthesis includes the type of glia involved in the corresponding neuronal paradigm, + designates that *orion* is involved in glia phagocytic action, - designates no *orion* function and N.D. non determined.

| Development (pruning or apoptosis) |  | injury |  |
| --- | --- | --- | --- |
| VNC synapses (astrocytes) | + | N.D |  |
| vCrz (astrocytes and cortex glia) | + | N.D |  |
| PDF-Tri (ensheathing glia and cortex) | + | N.D |  |
| N.D |  | Wing (L1 axons) (wrapping glia) | - |
| N.D |  | ORN (ensheathing glia) | - |
| NMJ (wrapping glia) | - | N.D |  |

### Drpr as an Orion receptor candidate

Interestingly, it was recently proposed that in larval da neurites Orion bridges phosphatidylserine exposed by neuron to Drpr receptor-expressing phagocytic epithelial cells during injury (Ji et al. 2022), indicating that Orion might be able to bind Drpr in this model. Furthermore, each neuron-glia phagocytosis model that we have analyzed in this study has been previously shown to be primarily driven by Drpr. Thus, *drpr* has a role in VNC synapse elimination and vCrz cell body removal (Tasdemir-Yilmaz and Freeman 2014). Interestingly, *drpr* does not seem involved in the elimination of vCrz^+^ neurites. In this particular case, the Crk/ Mbc/Ced-12 complex has been reported to be active, *via* an unknown receptor or signal, in glia to induce vCrz+ neurite elimination. This data point to Drpr and to an unknown receptor that might act *via* the Crk/ Mbc/Ced-12 complex, as major candidate receptors for Orion to activate glia phagocytosis during VNC development. Concerning central PDF-Tri neurons, it was recently reported that both their cell bodies and neurites require Drpr-dependent glial clearance (Vita et al. 2021). These data let to think that Orion might act *via* Drpr to eliminate the entire PDF-Tri neuronal tree. Olfactory nerve axotomy also triggers upregulation of Drpr in local ensheathing glial cells and consequently, in the absence of *drpr*, ensheathing glia fails to react and axonal debris lingers in the brain. Similarly, Drpr-dependent signaling mechanisms are required for efficient clearance of axonal debris in the L1 wing vein after wing injury (Neukomm et al. 2014). However, Orion does not seem to be required in any of those two injury paradigms. One possibility could be that Orion activates the Drpr pathway only during developmental remodeling and larval injury and therefore other ligands activate Drpr during adult neuron injury. In accordance with this idea, several ligands have been proposed as mediators of the OR neuron-ensheating glia crosstalk after ORN axotomy, like the Insulin-like peptides (Ilps) that belong to a class of injury-released factors (Musashe et al. 2016). Ilps are packaged into dense core vesicles (DCV) for release from injured axons shortly after injury.

They bind to the InR (*Insulin receptor*) in ensheathing glia and this in turn activates glial phagocytic activity *via* Drpr to promote clearance of degenerating axons. Ligands allowing communication between injured L1 nerve neurons and glial Drpr have not yet been described in the wing injury system. Furthermore, exploring the glia signaling pathways activated by Orion during neuronal degeneration in development and injury is an ongoing challenge to better understand mechanisms of neuron glia communication.

## Acknowledgements

We thank Jae H. Park for *Crz-GAL4* and *UAS-Casor* stocks, Marc Fremann for the *alrm-GAL4* lines, Jan-Adrianus Veenstra for the anti-Crz antibody, the Bloomington *Drosophila* Stock Center and VDRC for fly stocks, the BioCampus RAM-*Drosophila* facility (Montpellier, France), the imaging facility MRI, which is part of the UMS BioCampus Montpellier and a member of the National Infrastructure France-BioImaging, for help in confocal and image analysis and processing. The 1D4 anti-Fasciclin II hybridoma and the 8D12 anti-Repo monoclonal antibody developed by Corey Goodman were obtained from the Developmental Studies Hybridoma Bank, created by the NICHD of the NIH and maintained at The University of Iowa, Department of Biology, Iowa City, IA 52242. LJN was supported by a Swiss National Science Foundation SNSF Assistant Professor awards (176855 and 211015), the International Foundation for Research in Paraplegia (P180), and SNSF Spark (190919). C.P. was supported by grants from the INSB at the CNRS and from the ANR. Work in the laboratory of J.-M.D. was supported by the Centre National de la Recherche Scientifique, the Fondation pour la Recherche Médicale (Equipes FRM 2016) and the Agence National de la Recherche (ANR-ORIO).

