## Supplemental data for "Neuron secreted chemokine-like Orion is involved in the transformation of glial cells into phagocytes in different neuronal remodeling paradigms"

**Supplemental figure legends**

**Supplemental Fig 1. Orion is required for the elimination of vCrz$^+$ cell bodies and axons.** **(A-J)** Confocal z-stacks showing vCrz$^+$ neurons visualized by the expression of *Crz-GAL4* driven *UAS-mCD8-GFP* (green) at the indicated developmental stages in control **(A-E)** and *orion$^1$* mutants **(F-J)**. Horizontal axons were labeled by arrows, medial and lateral axons were labeled by arrowheads and an asterisk respectively in A. **(K)** Quantification of the number of vCrz$^+$ cell bodies, **(L)** vCrz$^+$ horizontal axons, **(M)** vCrz$^+$ medial axons and **(N)** vCrz$^+$ lateral axons in controls and *orion* mutants at 4 h and 6 h APF. Genotypes are listed in Supplementary list of fly strains. Error bars represent ± SEM (Mann-Whitney *U* test). Scale bars is 70 µm.

**Supplemental Fig 2. The elimination of vCrz$^+$ neurites and cell bodies does not seem mediated by ensheathing glia.** **(A, B)** Ensheathing glia visualized by the expression of *MZ0709-GAL4* driven *UAS-mCD8-GFP* (green) and vCrz$^+$ neurons by an anti-Crz antibody (red) at 4 h APF. **(B)** Arrows point to vCrz$^+$ cell bodies, which are not reached by ensheathing glia extensions and arrowhead to unengulfed neurites. See inset for higher magnifications of B in **B'** and **B''**. **A** is a z-projection confocal image, **B** is a single confocal plan. Genotypes are listed in Supplementary list of fly strains. Scale bar is 30 µm.

**Supplemental Fig 3. Nuclear targeting of Casor probe in the vCrz neurons.** Confocal z-stacks showing vCrz cell bodies visualized by the expression of *Crz-GAL4* driven *UAS-Casor* probe (green) **(A, B)** Intact casor probe monitored by intrinsic GFP was detected in most of the soma of controls **(A)** and *orion$^{\Delta C}$* **(B)** at 0 h APF (arrows). **(C, D)** Cleaved Casor probe was detected mainly in the nuclei in controls **(C)** and *orion$^{\Delta C}$* **(D)** new born flies (arrowheads) at 2 h APF. Scale bar is 40 µm **(E)** Percentage of vCrz$^+$ cell bodies containing Casor-GFP fluorescence in the nucleus, cytoplasm or in both nucleus and cytoplasm (nucleus + cytoplasm) at 0 and 2 h APF. The number of cell bodies is included in a parenthesis for each condition.

**Supplemental Fig 4. Orion mutant retains developmentally transient PDF-Tri neurons.** **(A-D)** Confocal z-stacks showing PDF **(A, B)** and PDF-Tri **(C, D)** neurons visualized by the expression of *PDF-GAL4* driven *UAS-mCD8-GFP* (green) and labeled with anti-PDF antibody (red) at the indicated time points in wild type **(A, C)** and $orion^1$ mutants **(B, D).** Note that the whole PDF neuron network is seen in A and B (asterisk in A) and that no differences are observed between controls and mutants in PDF regions other than the PDF-Tri one. Genotypes are listed in Supplementary list of fly strains. Scale bar is 50 µm. **(E)** Surfaces (in $\mu m^2$) occupied by the PDF-Tri arborization at 3 weeks in controls and *orion* mutants. Error bars represent ± SEM (Mann-Whitney *U* test). The number of brains analyzed is included in a parenthesis for each condition.

**Supplemental Fig 5. Astrocytes surrounding PDF-Tri neurons do not seem to eliminate PDF-Tri neurites. (A, E)** Confocal z-stacks showing astrocytic glia, PDF-Tri neurons and nuclei respectively visualized by the expression of *alrm-GAL4* driven *UAS-mCD8-GFP* (green), an anti-PDF antibody (red) and DAPI (blue) at 0 **(A)** and two **(E)** days. Different regions of the brain contained in these z-stacks are shown as single confocal plans (**B, C, F)** (SEZ region), (**D)** (MDBL region). Rectangles border regions showing PDF-Tri dendrites close to astrocytic glia, but engulfment is not observed. These regions are shown at higher magnification in **B'**, **C'**, **D'** and **F'**. Note the long and thin structure of the filamentous processes of the astrocytes in 2 days flies (arrows) that does not engulf neurites. Scale bars are 50 µm (A-D) and 30 µm (E-F).

**Supplemental Fig 6. Nuclear targeting of Casor probe in PDF-Tri neurons.** Confocal z-stacks showing PDF-Tri cell bodies visualized by the expression of *PDF-Tri-GAL4* driven *UAS-Casor* probe (green) and anti-PDF antibody co-staining (red) **(A, B)** Intact Casor probe monitored by intrinsic GFP was detected in most of the soma of controls **(A)** and $orion^{\Delta C}$ **(B),** at late pupal stage **(C, D)** Cleaved Casor probe was detected in most of the nuclei of controls **(C)** and $orion^{\Delta C}$ **(D)** new born (0 days) flies (arrowheads). Genotypes are listed in Supplementary list of fly strains. Scale bar is 30 µm **(E)** Representation of the percentage of PDF-Tri cell bodies containing Casor-GFP fluorescence in the nuclei, cytoplasm or both nuclei and cytoplasm (nuclei + cytoplasm) in pharates and new born flies. The number of cell bodies is included in a parenthesis for each condition.

**Supplemental Fig 7. Orion is not involved in the elimination of ORN debris by ensheathing glia after injury by palp ablation. (A, B)** Confocal z-stacks showing ORNs before injury (control) visualized by the expression of *Or85e-GAL4* driven *UAS-mCD8-GFP* (green) in wild type **(A)** and $orion^{1}$ mutants **(B). (C-G)** Confocal z-stacks showing ORNs 24 h after palp ablation visualized by the expression of *Or85e-GAL4* driven *UAS-mCD8-GFP* (green) in wild type **(C)** and $orion^{1}$ mutants **(D)** and 3 days after palp ablation in wild type **(F)** and $orion^{1}$ mutants **(G)**. Note the axon discontinuities observed in both conditions in C, D, 24 h after palp ablation (arrows in C and D) and that only disseminated debris are observed in both conditions 3 days after palp ablation. Genotypes are listed in Supplementary list of fly strains. Scale bar is 30 µm. n ≥ 20 brains for each condition.

**Supplemental Fig 8. NMJ dismantling during *Drosophila* metamorphosis is similar in controls and *orion* mutants. A-L**, Composite confocal z-stack images of muscle 4 NMJ in wild type larvae (L3) **(A-F)** and 6 h APF **(G-L)**, labeled with anti-DLG antibody (green) and anti-HRP antibody (red). At L3, the anti-DLG staining surrounds the bouton membranes as a thin layer in controls **(A-C)** and *orion* mutants **(D-F)**. **(G-L)** At 6 h APF the NMJ is similarly disorganized in both, controls **(G-I)** and orion mutants **(J-L)**, a high number of filopodia (arrow in H and K) and HRP-labelled debris were observed (arrowhead in K, L), the anti-DLG staining became discontinuous and fragmented. Individual synaptic boutons are no longer observed with the HRP-staining. The motor neuron became shorter and swollen at the synapse. Genotypes are listed in Supplementary list of fly strains. Scale bar is 20 μm. n ≥ 20 for each condition.

### Supplementary List of Fly Strains:

#### Figure 1:

**(A) *alrm-GAL4-GFP:*** *y* $w^{67c23}$ / *Y*; *alrm-GAL4 2x UAS-mCD8GFP/+*
**(B) *orion*$^{\Delta C}$*; alrm-GAL4-GFP:*** $w^{67c23}$ *orion*$^{\Delta C}$ / *Y*; *alrm-GAL4 2x UAS-mCD8GFP/+*
**(C, D) *control:*** *y* $w^{67c23}$ $sn^3$ *FRT19A* / Y or *y* $w^{67c23}$ $sn^3$ *FRT19A* / *y* $w^{67c23}$ $sn^3$ *FRT19A*
**(D, F) *orion*$^1$*:*** *y* $w^{67c23}$ $sn^3$ *orion* $^1$ *FRT19A* / Y or *y* $w^{67c23}$ $sn^3$ *orion* $^1$ *FRT19A* / *y* $w^{67c23}$ $sn^3$ *orion*$^1$ *FRT19A*

#### Figure 2:

**(A, F, H) *alrm-GAL4-GFP:*** *y* $w^{67c23}$ / *Y*; *alrm-GAL4 2x UAS-mCD8GFP/+*
**(B) *orion*$^{\Delta C}$*; alrm-GAL4-GFP:*** $w^{67c23}$ *orion*$^{\Delta C}$ / *Y*; *alrm-GAL4 2x UAS-mCD8GFP/+*
**(C) *control:*** *y* $w^{67c23}$ / *Y*; *alrm-GAL4 2x UAS-mCD8GFP/+;* ***orion*$^{\Delta C}$*:*** $w^{67c23}$ *orion*$^{\Delta C}$ / *Y*; *alrm-GAL4 2x UAS-mCD8GFP/+*
**(D) *alrm-GAL4-GFP:*** *w* / Y; alrm-GAL4 UAS-mCD8GFP hs-repo-FLP* $^{6.2}$*; FRT2A/FRT2A, TubGAL80*
**(E) *orion*$^{\Delta C}$*; alrm-GAL4-GFP:*** $w^{67c23}$ *orion*$^{\Delta C}$ */ Y; alrm-GAL4 UAS-mCD8GFP hs-repo-FLP* $^{6.2}$*; FRT2A/FRT2A, TubGAL80*
**(G, I) *orion*$^1$*; alrm-GAL4-GFP:*** *y* $w^{67c23}$ $sn^3$ *orion* $^1$ *FRT19A* / *Y*; *alrm-GAL4 2x UAS-mCD8GFP*

#### Figure 3:

**(A-D) *control :*** *y* $w^{67c23}$ */Y* or *y* $w^{67c23}$ / *y* $w^{67c23}$
**(E-H) *orion*$^1$ *:*** *y* $w^{67c23}$ $sn^3$ *orion* $^1$ *FRT19A* / *Y* or *y* $w^{67c23}$ $sn^3$ *orion* $^1$ *FRT19A*/ *y* $w^{67c23}$ $sn^3$ *orion* $^1$ *FRT19A*
**(I, J) *control:*** *y* $w^{67c23}$ */Y* or *y* $w^{67c23}$ / *y* $w^{67c23}$, ***orion*$^1$*:*** *y* $w^{67c23}$ $sn^3$ *orion* $^1$ *FRT19A* / *Y* or *y* $w^{67c23}$ $sn^3$ *orion* $^1$ *FRT19A*/ *y* $w^{67c23}$ $sn^3$ *orion* $^1$ *FRT19A*
**(K) *elav-GAL4; UAS-orion:*** *elav-GAL4 w UAS-CD8-GFP hs-FLP / Y or elav-GAL4 w UAS-CD8-GFP hs-FLP / y* $w^*$*; UAS-orion-B-myc / +*
**(L) *orion*$^{\Delta C}$ *elav-GAL4 :*** *elav-GAL4* $w^*$ *UAS-CD8-GFP orion*$^{\Delta C}$ / *Y*

**(M) *orion$^{\Delta C}$ elav-GAL4 ; UAS-orion:*** *elav-GAL4 $w^{*}$ UAS-CD8-GFP orion$^{\Delta C}$ ; UAS-orion-B-myc / +*
**(N) *orion$^{\Delta C}$ elav-GAL4 :*** *elav-GAL4 $w^{*}$ UAS-CD8-GFP orion$^{\Delta C}$ / Y*, ***elav-GAL4; UAS-orion:*** *elav-GAL4 w UAS-CD8-GFP hs-FLP / Y or elav-GAL4 w UAS-CD8-GFP hs-FLP / y $w^{*}$; UAS-orion-B-myc / +*, ***orion$^{\Delta C}$, elav-GAL4 :*** *elav-GAL4 $w^{*}$ UAS-CD8-GFP orion$^{\Delta C}$ / Y*

**(O) *repo-GAL4-GFP:*** *y $w^{67c23}$ $sn^{3}$ FRT19A / Y; repo-GAL4 UAS-mCD8GFP/+*
***(P) control :*** *y $w^{67c23}$ $sn^{3}$ FRT19A / Y; repo-GAL4 UAS-mCD8GFP/+* , ***orion$^{1}$:*** *y $w^{67c23}$ $sn^{3}$ orion $^{1}$ FRT19A/ Y; repo-GAL4 UAS-mCD8GFP/+*

**Figure 4 :**

**(A-C) *alrm-GAL4-GFP:*** *y $w^{67c23}$ / Y* or *$yw^{67c23}$ / $yw^{67c23}$*; *alrm-GAL4 2x UAS-mCD8GFP / Tb*
**(D, E) *NP2222-GAL4-GFP:*** *$w^{*}$ / Y*; or *$w^{*}$ / y $w^{67c23}$; NP2222-GAL4 / + ; 2x UAS-mCD8GFP/+*

**Figure 5:**

**(A, E, I) *PDF-GAL4-GFP:*** *y $w^{67c23}$ $sn^{3}$ FRT19A / Y; Pdf-GAL4 / 2 x UAS-mCD8-GFP; Pdf-GAL4 / +*
**(B, F, J) *orion$^{1}$; PDF-GAL4-GFP:*** *y $w^{67c23}$ $sn^{3}$ orion $^{1}$ FRT19A / Y; Pdf-GAL4 / 2 x UAS-mCD8-GFP; Pdf-GAL4 / +*
***(C, D, G, H, K, L) control:*** *y $w^{67c23}$ $sn^{3}$ FRT19A / Y; Pdf-GAL4 / 2 x UAS-mCD8-GFP; Pdf-GAL4 / +*, ***orion$^{1}$:*** *y $w^{67c23}$ $sn^{3}$ orion $^{1}$ FRT19A / Y; Pdf-GAL4 / 2 x UAS-mCD8-GFP; Pdf-GAL4 / +*
**(M) orion $^{\Delta C}$; UAS-orion :** *$w^{67c23}$ orion$^{\Delta C}$ / Y ; UAS-orion-B-myc / +.*
**(N) orion$^{\Delta C}$, elav-GAL4 ; UAS-orion :** *elav-GAL4 $w^{*}$ UAS-CD8-GFP orion$^{\Delta C}$ / Y ; UAS-orion-B-myc / +*
**(O, P) control, *:*** *y $w^{67c23}$ $sn^{3}$ FRT19A / Y; Pdf-GAL4 / 2 x UAS-mCD8-GFP; Pdf-GAL4 / +*, **orion $^{\Delta C}$; UAS-orion :** *$w^{67c23}$ orion$^{\Delta C}$ / Y ; UAS-orion-B-myc / +* **orion$^{\Delta C}$ , elav-GAL4 ; UAS-orion :** *elav-GAL4 $w^{*}$ UAS-CD8-GFP orion$^{\Delta C}$ / Y ; UAS-orion-B-myc / +*

**Figure 6:**

**(A-F') *MZ0709-GAL4-GFP:*** *w $^{1118}$ / Y*; or *w $^{1118}$ / y $w^{67c23}$; MZ0709-GAL4 / +; 2x UAS-mCD8GFP/+*
**(G-I) *NP2222-GAL4-GFP:*** *$w^{11118}$ / Y*; or *$w^{1118}$ / y $w^{67c23}$; NP2222-GAL4 / +; 2x UAS-mCD8GFP/+*

**Figure 7:**

**(A, A') wild type:** *y, $w^{67c23}$, $sn^{3}$, FRT19A / $w^{-}$; dpr1–GAL4, 5xUAS–mCD8-GFP, ase-FLP$^{2b}$ / +; FRT2A/tub–GAL80, FRT2A*
**(B, B') orion$^{1}$/+:** *y, $w^{67c23}$, $sn^{3}$, orion$^{1}$, FRT19A / $w^{-}$; dpr1–GAL4, 5xUAS–mCD8-GFP, ase-FLP$^{2b}$ / +; FRT2A/tub–GAL80, FRT2A*
**(C, C') wild type:** *y, $w^{67c23}$, $sn^{3}$, FRT19A / Y; dpr1–GAL4, 5xUAS–mCD8-GFP, ase-FLP$^{2b}$ / +; FRT2A/tub–GAL80, FRT2A*

**(D, D') orion$^1$:** *y, $w^{67c23}$, $sn^3$, $orion^1$, FRT19A / Y; dpr1–GAL4, 5xUAS–mCD8-GFP, ase-$FLP^{2b}$ / + ; FRT2A/tub–GAL80,FRT2A*

**Supplementary Figure 1:**

**(A-E)** ***Crz-GAL4-GFP :*** *y $w^{67c23}$ $sn^{3\ 1}$ FRT19A / Y; + / 2 x UAS-mCD8-GFP; Crz-GAL4 / +*
**(F-J)** ***orion$^1$; Crz-GAL4-GFP:*** *y $w^{67c23}$ $sn^3$ orion $^1$ FRT19A / Y; + / 2 x UAS-mCD8-GFP; Crz-GAL4 / +*
**(K-N)** ***control:*** *y $w^{67c23}$ $sn^{3\ 1}$ FRT19A / Y; + / 2 x UAS-mCD8-GFP; Crz-GAL4 / + ;* ***orion$^1$:*** *: y $w^{67c23}$ $sn^3$ orion $^1$ FRT19A / Y; + / 2 x UAS-mCD8-GFP; Crz-GAL4 / +,* **yw:** *y $w^{67c23}$ ,* ***orion$^{\Delta C}$****: $w^{67c23}$ $orion^{\Delta C}$*

**Supplementary Figure 2:**

**(A-B'')** ***MZ0709-GAL4-GFP:*** *$w^{1118}$ / Y*; or *w $^{1118}$ / y $w^{67c23}$; MZ0709-GAL4 / +; 2x UAS-mCD8GFP/+*

**Supplementary Figure 3:**

**(A, C)** ***UAS-Casor; Crz-GAL4:*** *+ / UAS-Casor; Crz-GAL4 / +*
**(B, D*)*** ***orion$^{\Delta C}$; UAS-Casor; Crz-GAL4: :*** *$w^{67c23}$ $orion^{\Delta C}$; + / UAS-Casor; Crz-GAL4 / +*
**(E)** ***control:*** *+ / UAS-Casor; Crz-GAL4 / +* ***orion$^{\Delta C}$:*** *$w^{67c23}$ $orion^{\Delta C}$; + / UAS-Casor; Crz-GAL4 / +*

**Supplementary Figure 4:**

**(A, C)** ***PDF-GAL4-GFP:*** *y $w^{67c23}$ $sn^3$ FRT19A / Y; Pdf-GAL4 / 2 x UAS-mCD8-GFP; Pdf-GAL4 / +*
**(B, D)** ***orion$^1$; PDF-GAL4-GFP:*** *y $w^{67c23}$ $sn^3$ orion $^1$ FRT19A / Y; Pdf-GAL4 / 2 x UAS-mCD8-GFP; Pdf-GAL4 / +*
**(E)** ***control:*** *y $w^{67c23}$ $sn^3$ FRT19A / Y; Pdf-GAL4 / 2 x UAS-mCD8-GFP; Pdf-GAL4 / +,* ***orion$^1$:*** *y $w^{67c23}$ $sn^3$ orion $^1$ FRT19A / Y; Pdf-GAL4 / 2 x UAS-mCD8-GFP; Pdf-GAL4 / +*

**Supplementary Figure 5:**

**(A-F')** ***alrm-GAL4-GFP:*** *$yw^{67c23}$ / Y* or *$yw^{67c23}$ / $yw^{67c23}$*; *alrm-GAL4 2x UAS-mCD8GFP/ alrm-GAL4 2x UAS-mCD8GFP*

**Supplementary Figure 6:**

**(A, C)** ***UAS-Casor; PDF-GAL4:*** *yw* / Y; Pdf-GAL4 / UAS-Casor; Pdf-GAL4 / +*
**(B, D*)*** ***orion$^{\Delta C}$; UAS-Casor; PDF-GAL4*** *$w^{67c23}$ $orion^{\Delta C}$/ Y; Pdf-GAL4 / UAS-Casor; Pdf-GAL4 / +*

**(E)** ***control:*** *yw* / Y; Pdf-GAL4 / UAS-Casor; Pdf-GAL4 / +* ***orion***$^{\Delta C}$**:** *w*$^{67c23}$ *orion*$^{\Delta C}$*; Pdf-GAL4 / UAS-Casor; Pdf-GAL4 / +*

**Supplementary Figure 7:**

**(A, C, F) Or85e-GAL4-GFP:** *y w*$^{67c23}$ *sn*$^{3}$ *FRT19A / Y; 2 x UAS-mCD8GFP / +; Or85e-GAL4 / +*
**(B, D, G)** ***orion***$^{1}$***;*** **OR85e-GAL4-GFP:** *y w*$^{67c23}$ *sn*$^{3}$ *orion* $^{1}$ *FRT19A / Y; 2 x UAS-mCD8GFP / +; Or85e-GAL4 / +*

**Supplementary Figure 8:**

**(A-C, G-I)** ***control*****:** *y w*$^{67c23}$ *sn*$^{3}$ *FRT19A /* Y or *y w*$^{67c23}$ *sn*$^{3}$ *FRT19A / y w*$^{67c23}$ *sn*$^{3}$ *FRT19A*
**(D-F, J-L)** ***orion***$^{1}$***:*** *y w*$^{67c23}$ *sn*$^{3}$ *orion* $^{1}$ *FRT19A /* Y or *y w*$^{67c23}$ *sn*$^{3}$ *orion* $^{1}$ *FRT19A / y w*$^{67c23}$ *sn*$^{3}$ *orion*$^{1}$ *FRT19A*

Supplemental Fig. 1

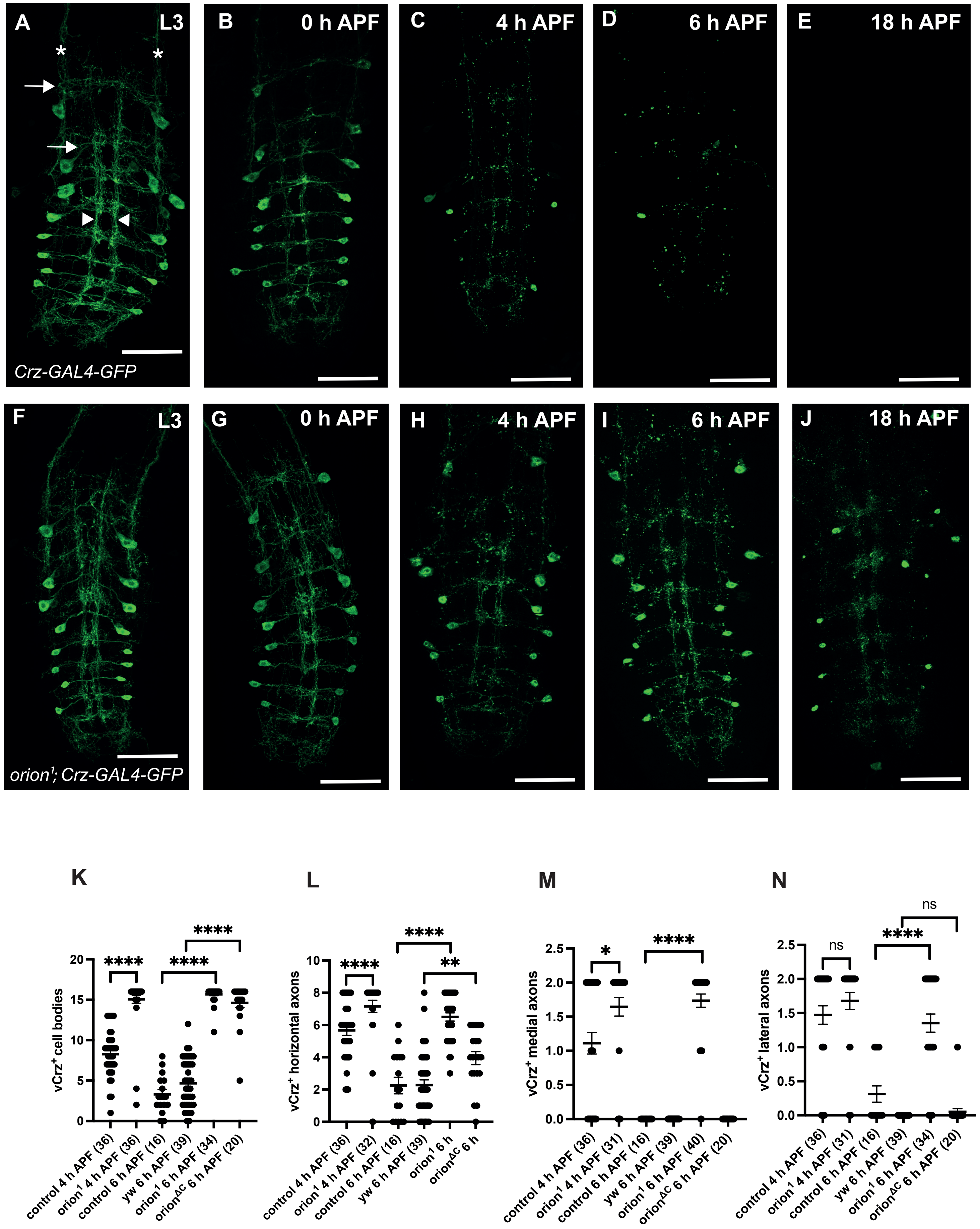

Supplemental Fig. 2

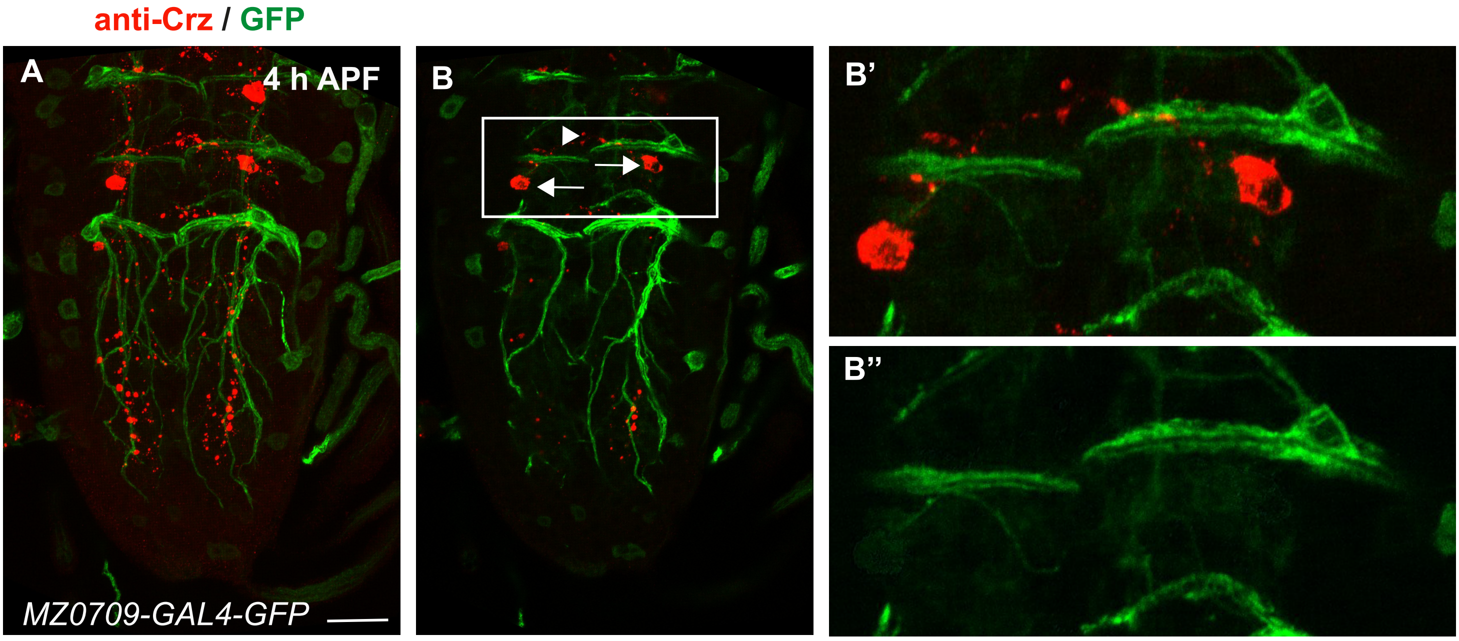

Supplemental Fig. 3

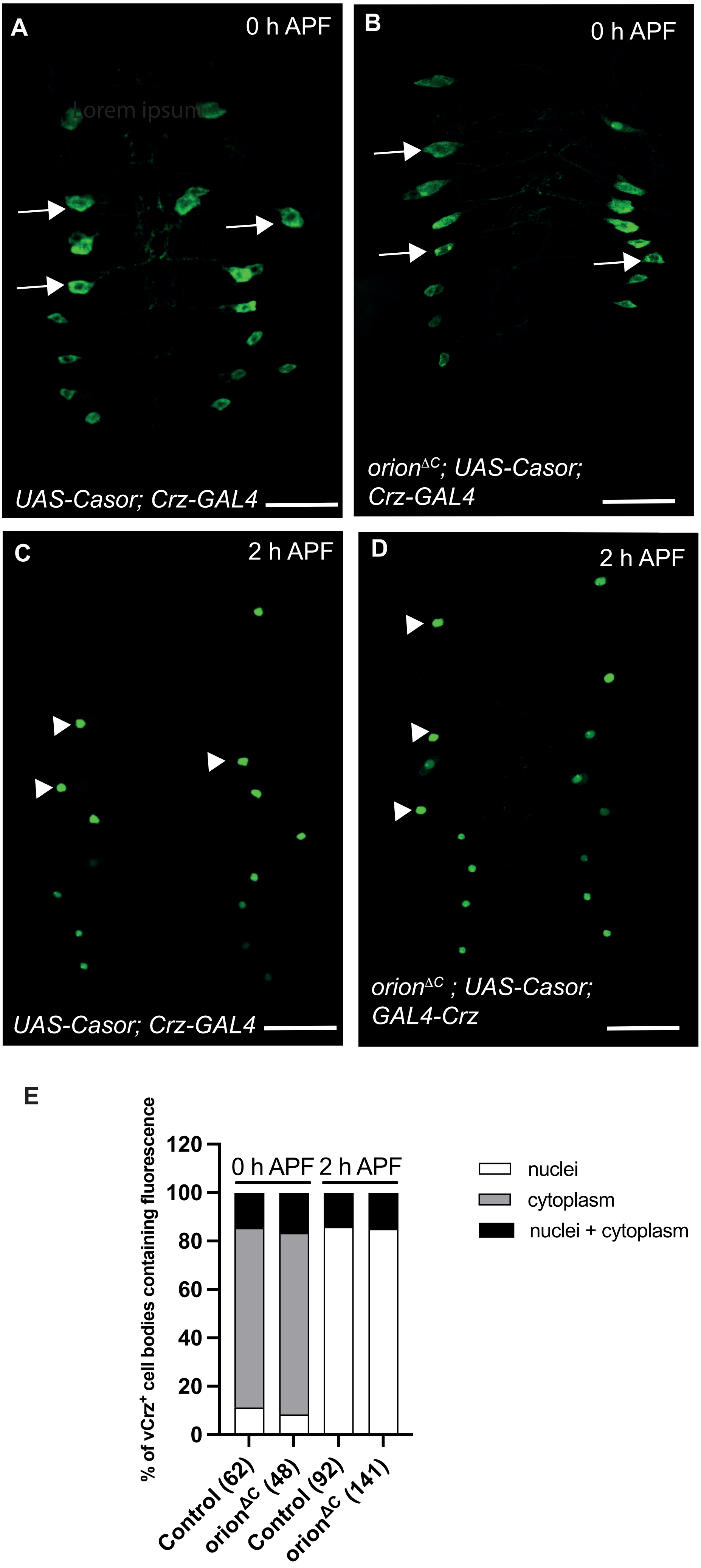

Supplemental Fig. 4

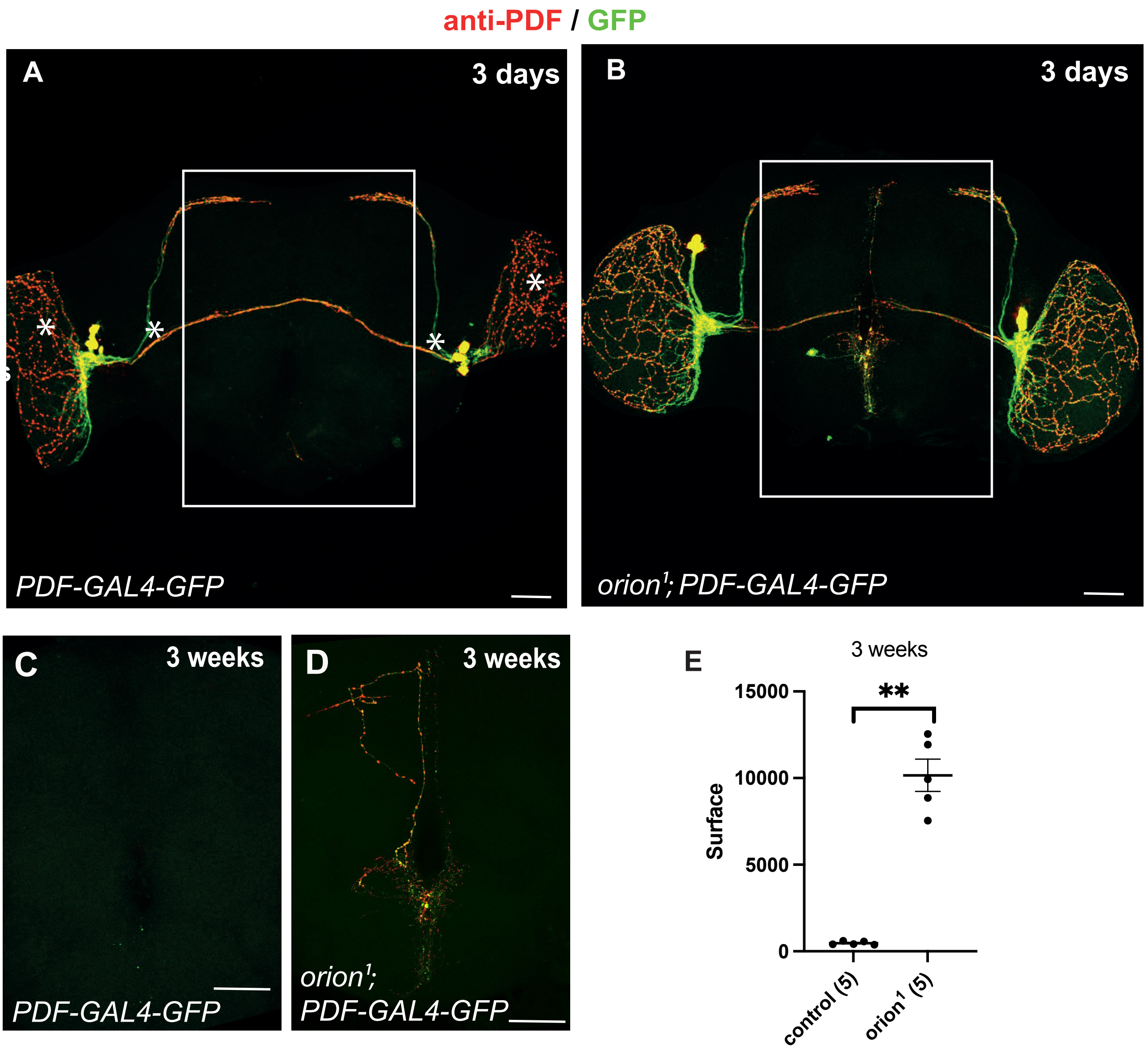

Supplemental Fig. 5

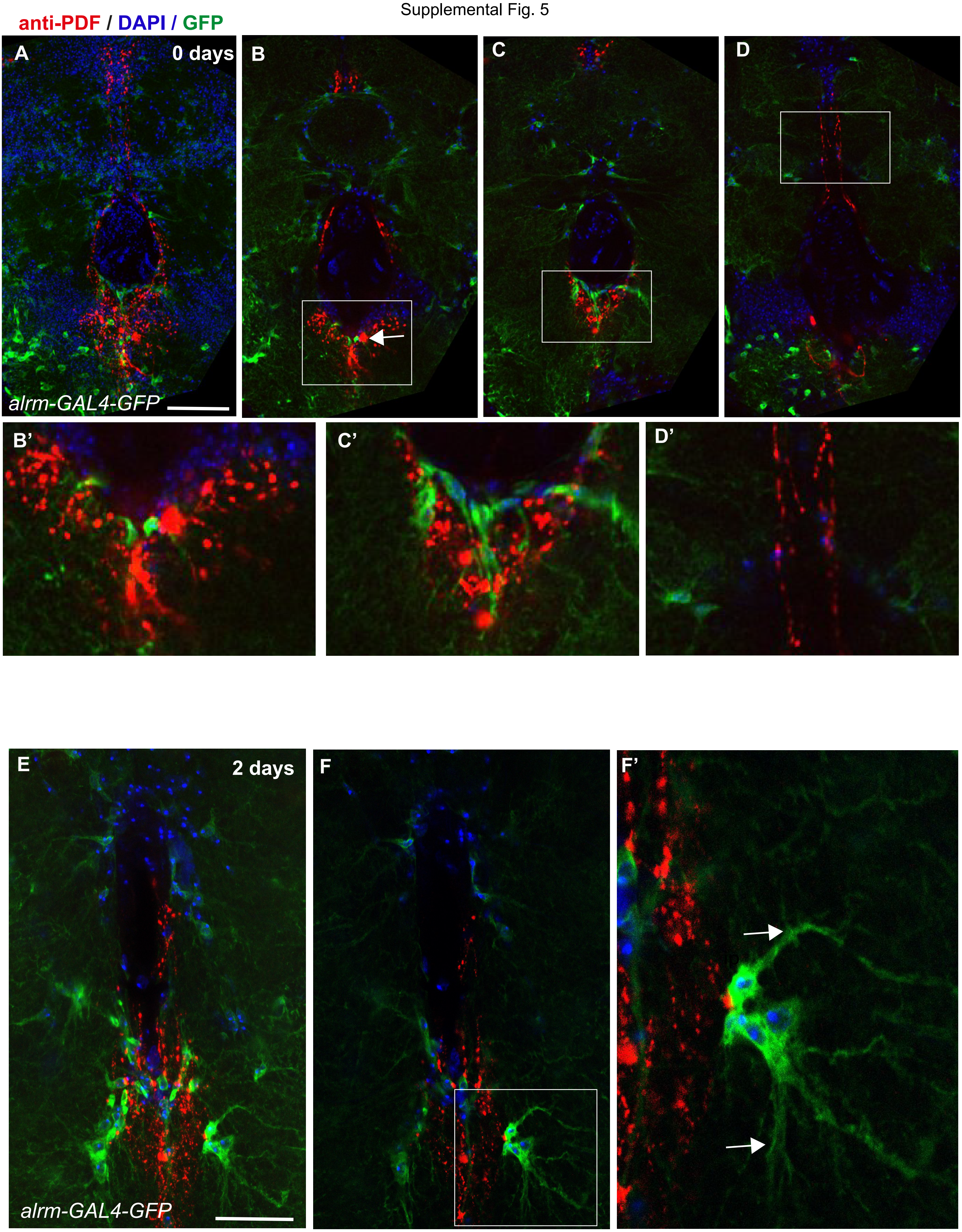

Supplemental Fig. 6

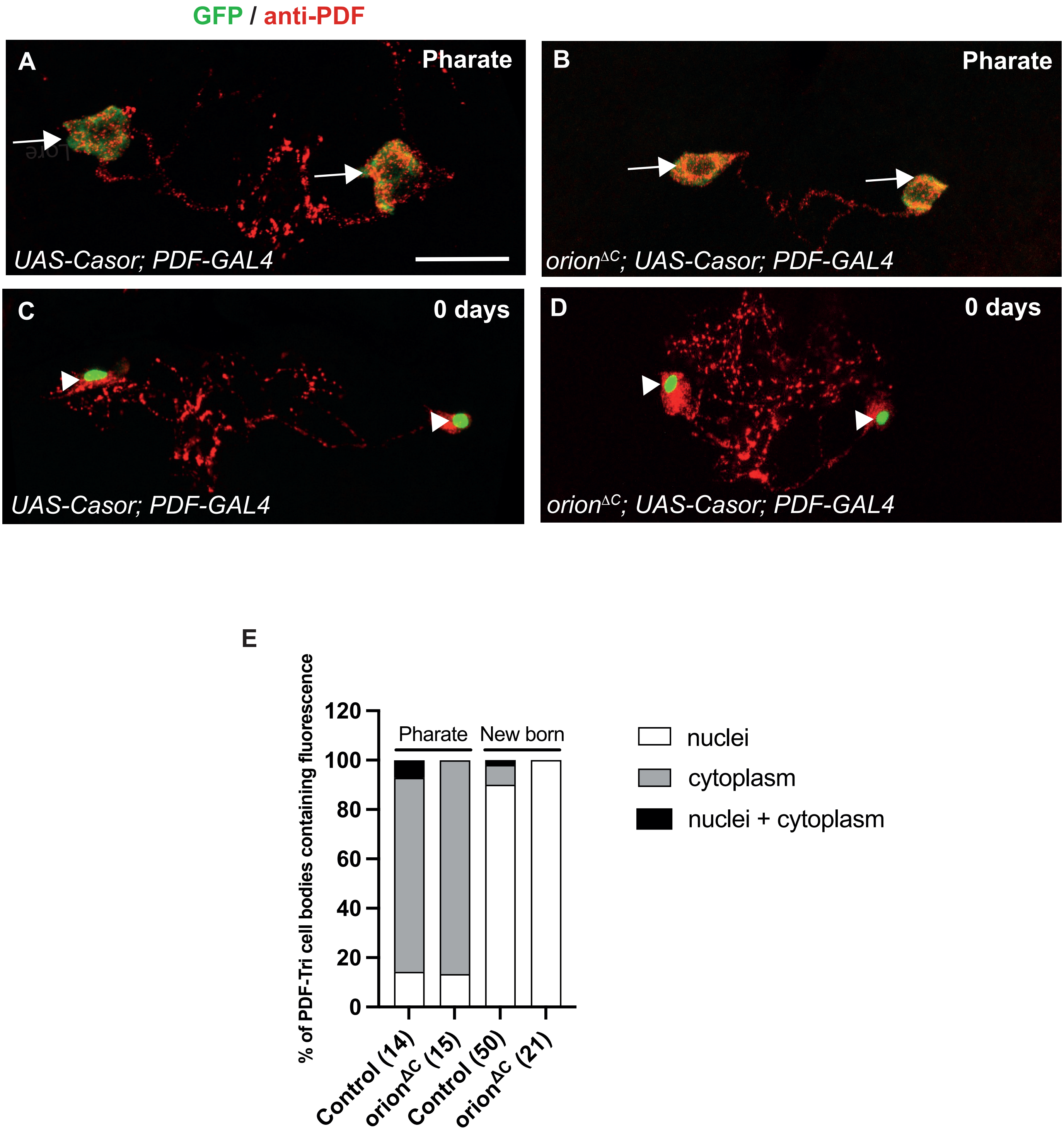

Supplemental Fig. 7

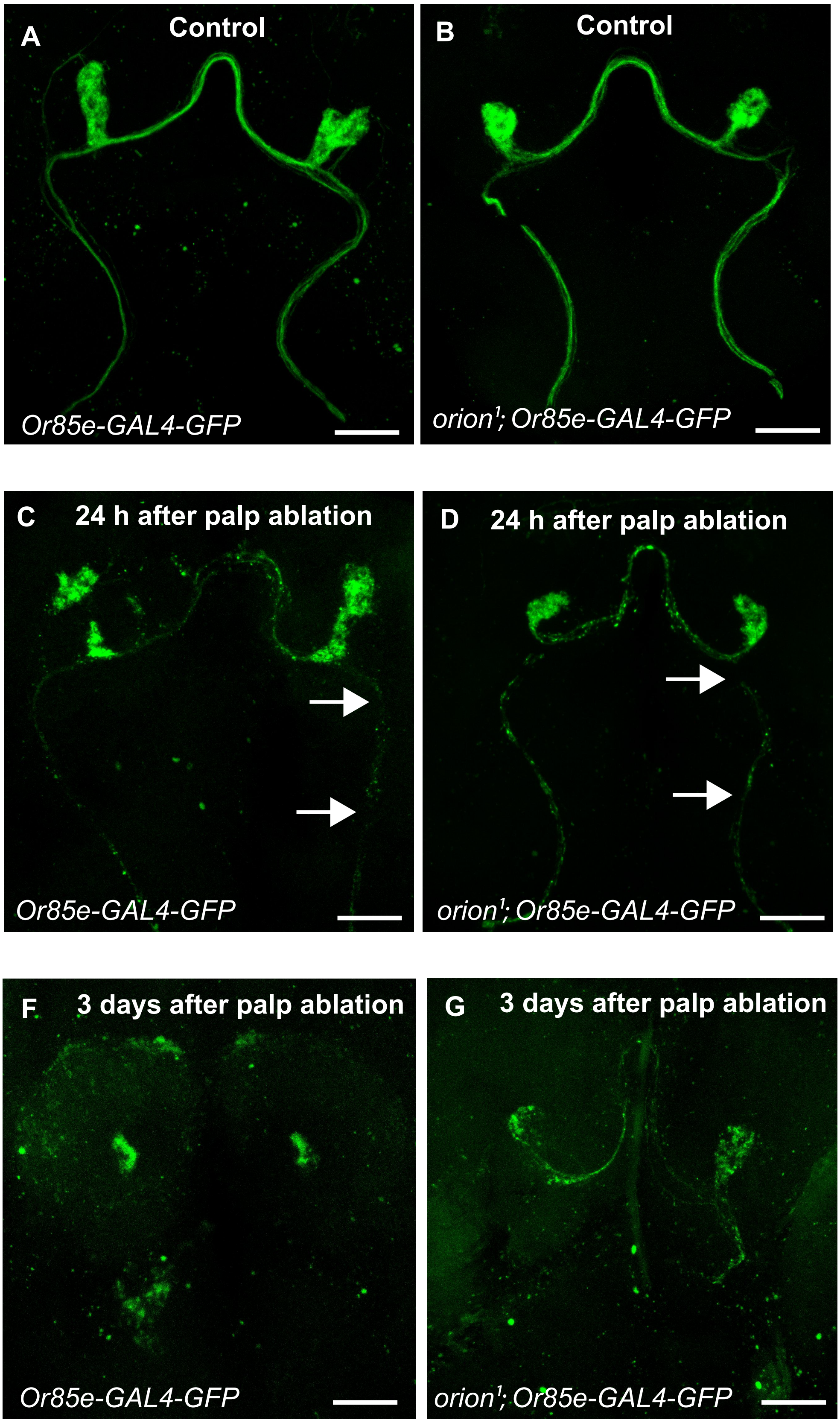

Supplemental Fig.8

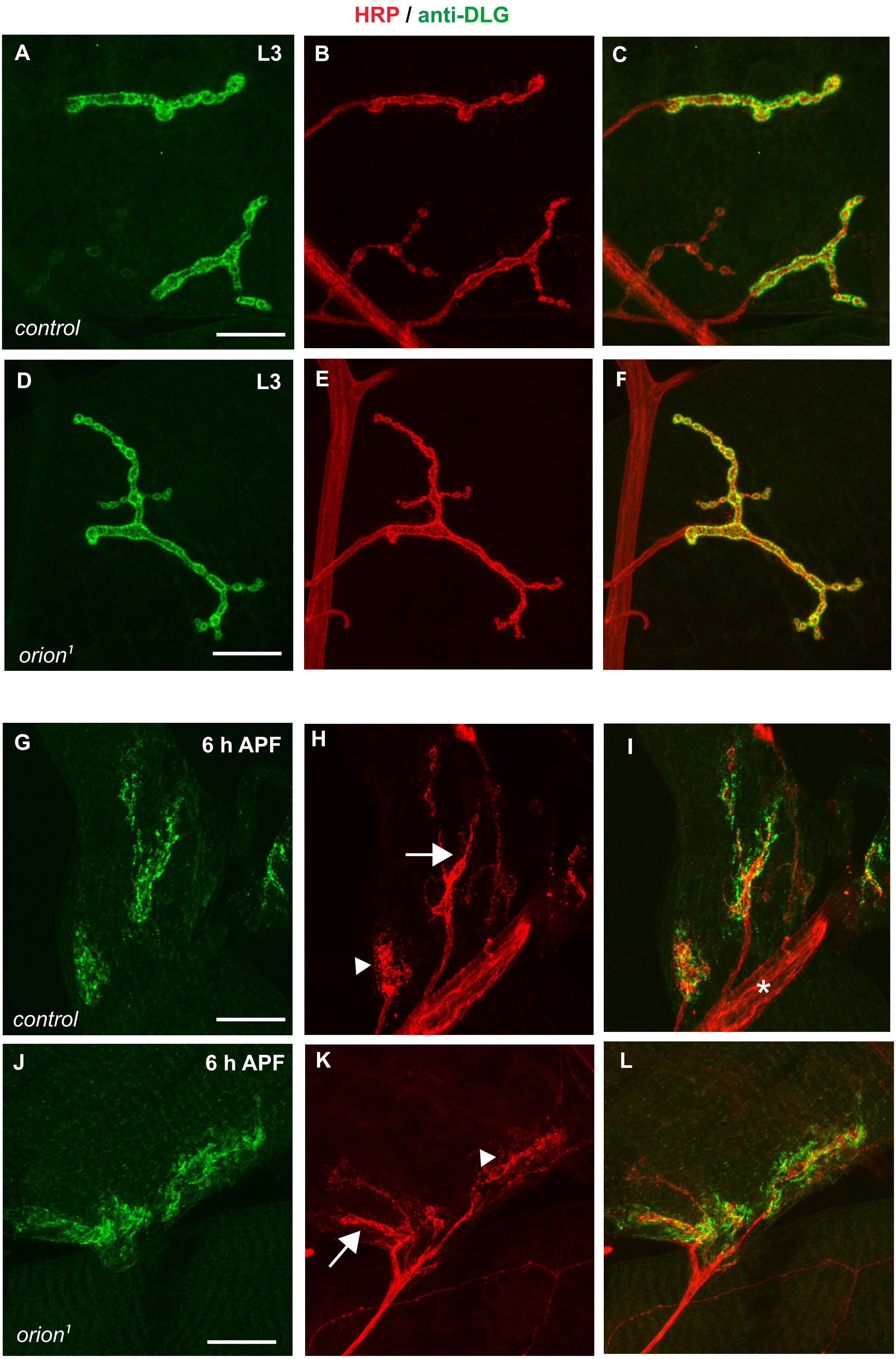
